# Human and mouse cerebellar inhibitory circuits in dystonic crisis and their modulation with therapeutic stimulation

**DOI:** 10.64898/2026.04.01.715774

**Authors:** Alejandro G. Rey Hipolito, Michael P. Dew, Jason S. Gill, Janelle E. Allen, Karissa A. Chesky, Mariam Hull, Roy V. Sillitoe

## Abstract

Dystonia is a neurological movement disorder characterized by abnormal muscle contractions that, at their most severe, lead to a life-threatening condition - dystonic crisis. Yet, therapeutic options remain limited by our incomplete understanding of what neural circuits underly the condition. Although cerebellar nuclei neurons are implicated in the origin of baseline dystonic symptoms, it is unclear whether their activity drives dystonic crisis. To explore this role, we found that cerebellar abnormalities and neural inhibition were recurring targets in patients with dystonic crisis, implicating cerebellar inhibitory neurons in its development. We devised a mouse genetics approach to test whether inhibitory cerebellar nuclei neurons (iCNNs) induce dystonic crisis. Directional optogenetic modulation of iCNNs induced dystonic crises on-demand and alleviated spontaneous crises in mice that mimic spontaneous dystonic crises. To investigate whether iCNNs interact with other motor areas during dystonic crisis, we identified monosynaptic iCNN projections to the centrolateral nucleus of the thalamus (CL). Deep brain stimulation of the CL alleviated dystonic crises induced by iCNN photoactivation. Our data uncover a cell type-specific cerebellar origin of dystonic crisis and highlight its therapeutic potential.

## Introduction

Dystonia is a neurological movement disorder characterized by sustained or intermittent muscle contractions that cause abnormal movements, postures, or both (*1*). In the most severe form of the condition, referred to as dystonic crisis (also termed dystonic storm or status dystonicus), the severity and persistence of dystonia require immediate hospitalization (*2*). Unfortunately, reaching this stage of devastating symptom intensity leads to mortality in up to 10-12.5% of patients, mostly comprising children with an existing diagnosis of dystonia (*3*, *4*). Relapses are common with a recurrence risk of 25% in the first year, and up to 38.6% at 27 months (*5*). Treatment options remain limited with variable success and include many complex pharmaceutical strategies or last-resort surgical interventions such as deep brain stimulation (DBS) or an intrathecal or intraventricular baclofen pump (*6*). Thus, there is an unmet need to develop additional therapeutic options for patients with dystonic crisis, which may be facilitated by identifying the exact neural circuits that drive the condition and targeting them with more precise and effective therapies.

It is unclear what specific neural circuits instigate dystonic crisis and how their dysfunction drives this debilitating condition. However, several lines of evidence suggest that inhibitory neurons and motor structures, especially the cerebellum, contribute to dystonic crisis. Ablation and DBS of motor structures including the globus pallidus and thalamus are therapeutically effective in a subset of patients in dystonic crisis (*7*), supporting a role for areas within the motor network in this condition. In dystonic crisis, treatments routinely involve drugs that target neural inhibition including benzodiazepines, alpha-2 adrenergic agonists, and anticholinergics (*3*, *5*, *6*, *8*), suggesting that inhibition in the central nervous system may play a role in the condition just as has been found in other forms of dystonia (*9–11*). Clinical studies have reported a high incidence of dystonic crisis in patients with cerebral palsy (*3*, *4*, *12*, *13*), a disorder in which cerebellar dysfunction is thought to play a prominent role (*14–16*), providing an indirect link to cerebellar dysfunction and dystonic crisis. However, these compelling connections require further investigation.

Although few studies on dystonic crisis exist, preclinical evidence in dystonia highlight a role for cerebellar nuclei neurons, which form the predominant output from the cerebellum to the rest of the motor network (*17*). In mice, lesioning the superior cerebellar peduncle, the cerebellar outflow tract through which cerebellar nuclei neurons project, was recently shown to cause severe dystonia similar to dystonic crisis (*18*), suggesting that cerebellar nuclei neurons may contribute to dystonic crisis. In animal studies of paroxysmal dystonia, dystonic episodes correlated with changes in cerebellar nuclei neuron activity. In one study, increased activity of inhibitory cerebellar nuclei neurons (iCNNs) correlated with the onset of dystonic episodes (*19*), suggesting that the cell type-specificity of cerebellar nuclei neurons may determine their contribution to dystonia. Little is known about the iCNN output pathway, though recent studies have described its termination throughout the brain (*20*, *21*), which may subserve unknown functions in health and disease. Accurately distinguishing the specific roles of the intermingled cerebellar nuclei neurons may be critical for developing better treatments for dystonia and other movement disorders, as was demonstrated for the basal ganglia in a mouse model of Parkinson’s disease (*22*).

To assess the role of iCNNs in dystonic crisis, we started by examining clinical data to study the possible contribution of the cerebellum and inhibitory neurons to dystonic crisis. To complement this clinical study, we sought to manipulate iCNNs in a mouse model of dystonia, and then test whether motor behaviors that resemble dystonic crisis arise. We therefore devised a conditional mouse genetic model to test the hypothesis that modulation of the iCNN output pathway induces dystonic crisis. Using a previously described animal model of dystonia, we genetically expressed opsins in *Ptf1a*-expressing neurons, including iCNNs, and optogenetically modulated them through their projections in the superior cerebellar peduncle. To examine if and how dystonic crises emerge, we combined this approach with behavioral assays and anatomical examination. We also tested whether iCNNs interact with another motor area implicated in dystonia, the centrolateral nucleus of the thalamus (CL), by using anatomical tracing. We then tested the hypothesis that DBS targeted to the CL would modulate the induced dystonic crises. Our data uncover a cerebellar origin of dystonic crisis and inspire the development of targeted therapies.

## Results

### Cerebellar abnormalities and inhibitory neural function are implicated in the incidence and treatment of dystonic crisis in human pediatric-onset dystonia

As previous clinical data only indirectly linked cerebellar function to dystonic crisis (*13*, *15*), we sought to identify more direct clinical evidence that cerebellar dysfunction leads to dystonic crisis and support further preclinical investigations into the precise cerebellar circuits that may be involved. Accordingly, we investigated the clinical characteristics (**Table 1**) of patients admitted with dystonic crisis (**<u>Supplementary Movie 1</u>**) at Texas Children’s Hospital in Houston, Texas between 2015 and 2019. Of the 49 patients admitted, 19 patients were defined as having dystonia associated with a genetic or progressive etiology while the remaining 30 patients (20 males, 10 females; between 3 months and 20 years of age at admission) had an underlying etiology associated with acquired injury (**Figure 1A and Table 1)**. Of the 30 patients with acquired injury, cerebellar abnormalities were observed in the MRI imaging of 18 (60%; 11 males, 7 females; between 3 months and 18 years of age at admission) patients while 12 of 30 patients (40%; 9 males, 3 females; between 3 and 20 years of age at admission) showed imaging with abnormalities exclusively in the supratentorial regions comprising the cerebral hemispheres, basal ganglia, thalamus, and other structures outside the cerebellum (**Figure 1B and Table 1**), suggesting that cerebellar abnormalities predispose a substantial number of patients to dystonic crisis. The range of cerebellar abnormalities found in patients with dystonic crisis encompassed focal cerebellar only to focal cerebellar with associated supratentorial to diffuse cerebellar injury (**Figure 1C**). When comparing the therapies that had clear clinical benefit (as determined by parents and clinicians) across the 30 patients with acquired injury, benzodiazepines and alpha-2 adrenergic agonists were the most widely used and successful (**Figure 1D**). Patients with cerebellar abnormalities responded preferentially to alpha-2 adrenergic agonists rather than benzodiazepines. Interestingly, benzodiazepines and alpha-2 adrenergic agonists have inhibitory effects on the central nervous system (*6*, *23*, *24*). In cerebellar nuclei neurons, the corresponding receptors for these two drugs have been identified (*25*, *26*), and alpha-2 adrenergic agonists have been found to modulate their GABA responses (*27*), indicating a potential role for GABAergic cerebellar nuclei neurons in the incidence and treatment of dystonic crisis. Patients with cerebellar abnormalities tended to have longer lengths of hospitalization (Unpaired t-test with Welch’s correction p=0.167, f=0.001) and days to improvement (Unpaired t-test with Welch’s correction p=0.104, f=0.004) following their admission with dystonic crisis (**Figure 1E**). To our knowledge, this is the first clinical evidence suggesting a direct link between cerebellar developmental abnormalities and dystonic crisis. Together, these results strongly support a role for the cerebellum and inhibitory neural circuitry in driving dystonic crisis in human pediatric-onset dystonia. As iCNNs represent the intersection of cerebellar function, neural inhibition, and cerebellar control of the motor network, we hypothesized that iCNNs instigate the mechanisms that trigger dystonic crisis.

**Figure 1:**
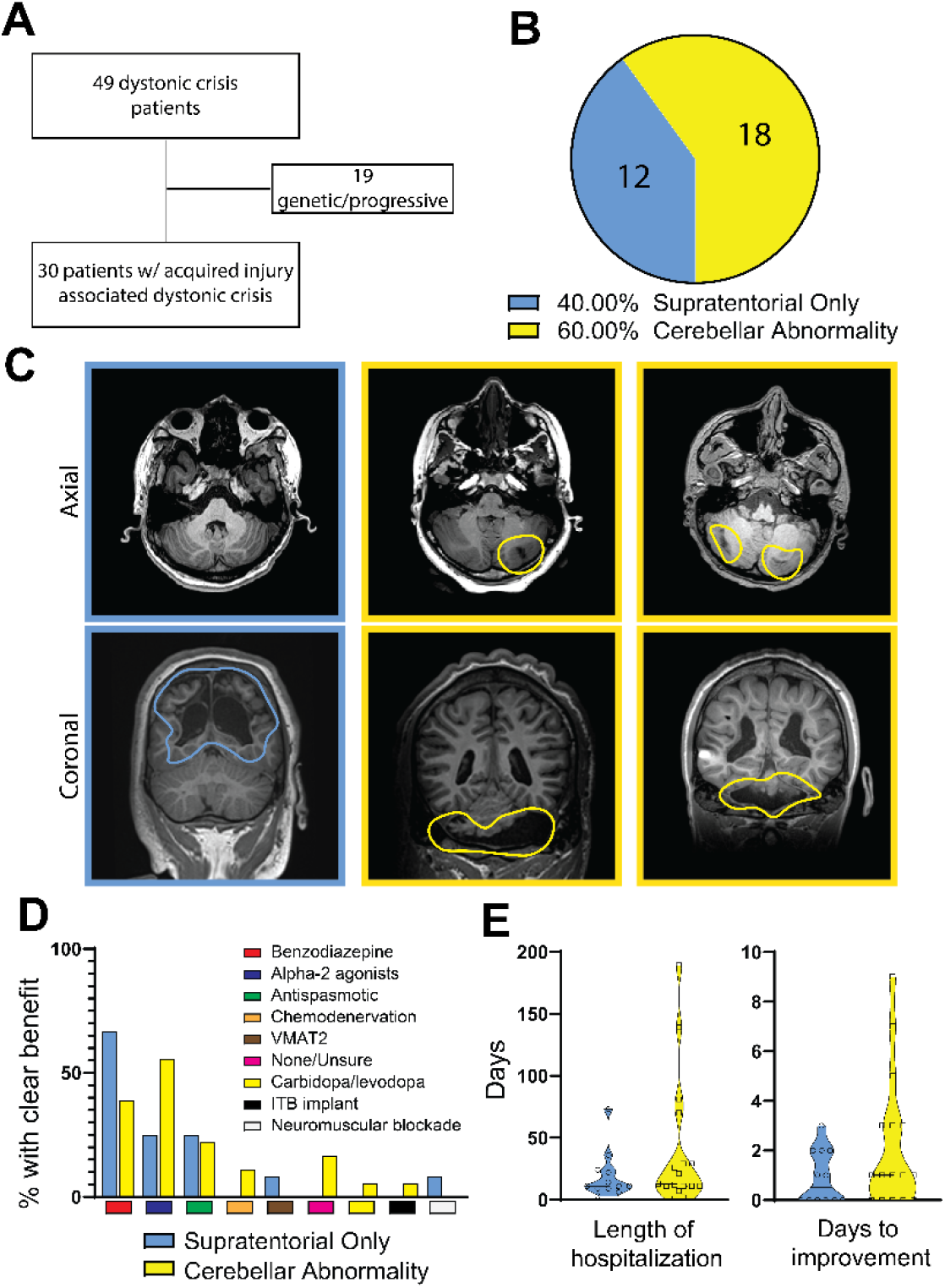
Cerebellar abnormalities and inhibitory neural function are implicated in the incidence and treatment of dystonic crisis. (**A**) Patient flow chart showing the division of 49 patients with dystonic crisis into groups based on the etiology of their dystonia: genetic/progressive (19 patients) or acquired injury (30 patients). (**B**) Percentage of patients admitted for dystonic crisis that had abnormalities in the cerebellum with or without supratentorial findings or only supratentorial abnormalities (blue) on imaging. (**C**) Examples of imaging in patients with supratentorial only injury (left column, outlined in blue) or cerebellar findings (middle and right column, outlined in yellow with abnormalities encircled). The top row depicts the axial plane through the cerebellum on T1 weighted imaging. The top left depicts the cerebellum at the level of the superior cerebellar peduncles, the top middle depicts a sequela of isolated left cerebellar hemispheric stroke, the top right depicts bilateral cerebellar hemispheric encephalomalacia (outlined in blue). The bottom row depicts the coronal plane through the cerebellum and the parietal lobe on T1 weighted imaging. The bottom right shows the normal cerebellum with extensive bilateral periventricular encephalomalacia. The bottom middle and right show patients with extensive bilateral cerebellar encephalomalacia. (**D**) Comparison of therapies that had clear clinical benefit across the 30 patients with acquired injury associated with dystonic crisis. Benzodiazepines and alpha-2 adrenergic agonists were the most widely used and successful. Patients with cerebellar abnormalities responded preferentially to alpha-2 adrenergic agonists rather than benzodiazepines. (**E**) Patients with cerebellar abnormalities tended to have longer lengths of hospitalization (Unpaired t-test with Welch’s correction p=0.167, f=0.001) and days to improvement (Unpaired t-test with Welch’s correction p=0.104, f=0.004) following their admission with dystonic crisis.

**Table 1.** Clinical characteristics of patients with dystonic crisis, also known as status dystonicus (SD). Table outlines clinical characteristics of 30 patients admitted to Texas Children’s Hospital from 2015-2019 with acquired etiologies of dystonic crisis. Top grouping includes patient with only supratentorial findings, while bottom grouping includes those with cerebellar findings. Summarized imaging findings are as indicated in the respective columns. Ages are shown in years. “Dystonia phenomenology” refers to the description of dystonia based on consensus classifications while “Description of phenomenology” indicates the patient specific presentation of dystonia during SD. Patients with imaging depicted in figure 1 are as follows: # = top left, $ = bottom left, € = top middle, % = bottom middle, £ = top right, & = bottom right. Abbreviations of treatments with clinical benefit: BZ = benzodiazepine; α-2 = Alpha-2 agonists; AS = antispasmodic agents; ChemD = chemodenervation; VMAT2 = VMAT-2 inhibitor; None = no therapy with clinical benefit; Dopa = dopaminergic agonism; ITB = intrathecal baclofen pump; NmB = neuromuscular blockade.

| Sex | Age of dystonia onset | Dystonia phenomenology | Etiology | Supratentorial Imaging Findings | Posterior Fossa Imaging Findings | Age of admission for status dystonicus | Description of phenomenology | BZ | α-2 | AS | ChemD | VMAT2 | None | Dopa | ITB | NmB | Days to improvement |
|---|---|---|---|---|---|---|---|---|---|---|---|---|---|---|---|---|---|
| F | 10 | generalized | Anoxic Brain Injury | Bilateral Cerebral encephalomalacia, thalamic gliosis, basal ganglia signal abnormalities |  | 20 | Intermittent stiffening and tremor | x |  |  |  |  |  |  |  |  |  |
| M | 0.25 | generalized | HIE | Basal ganlia signal abnormality, mild cerebral volume loss |  | 17 | Dystonia involving proximal portions of all extremites | x |  | x |  |  |  |  |  |  | 2 |
| M | 5 | generalized | Kernicterus | Mild signal change in basal ganglia |  | 11 | Dystonic and dyskinetic movements of all extremities | x |  |  |  |  |  |  |  |  | 0 |
| M | 15 | generalized | Prematurity | Severe bilateral cerebral encephalomalacia, thalamic volume loss. |  | 15 | Generalized dystonia | x | x |  |  |  |  |  |  |  | 0 |
| # M | 10 | generalized | Prematurity | abnormal signal in bilateral deep gray matter, periventricular leukomalacia. |  | 14 | Generalized severe increased tone |  | x |  |  |  |  |  |  |  | 0 |
| \$ F | 0.75 | generalized | Perinatal Injury | Bilateral occipitoparietal encephalomalacia and periventricular leukomalacia |  | 4 | Intermittent opithotonic posturing | x |  | x |  |  |  |  |  |  | 1 |
| M | 0.83 | generalized | HIE | Diffuse cortical and deep gray volume loss, thelamic calcifications. |  | 12 | Extensor posturing |  |  |  |  |  |  |  |  |  | 0 |
| M | 1 | generalized | HIE | Mild cerebral atrophy with severe arophy of bilateral basal ganglia and thalami. |  | 5 | Not characterized | x |  |  |  |  |  |  |  |  | 0 |
| M | 3 | generalized | NMDA Encephalitis | Basal ganglia encephalomalacia |  | 3 | Posturing of all extremities |  |  |  |  | x |  |  |  | x | 1 |
| M | 0.17 | generalized | HIE | Cerebral volume loss and abnormal singal of bilateral basal ganglia and thalami. |  | 10 | Mixed spastic/dystonic posturing | x |  |  |  |  |  |  |  |  | 2 |
| M | 1 | generalized | Kernicterus | Cerebral volume loss. |  | 10 | Generalized dystonia |  | x |  |  |  |  |  |  |  | 2 |
| F | 14 | generalized | Anoxic Brain Injury | Diffuse cerebral, basal ganlia, and thalamic edema. |  | 15 | UE>LE without spontaneous movements | x |  | x |  |  |  |  |  |  | 0 |
| M | 4 | generalized | Anoxic Brain Injury | Cerebral white matter and thalamic volume loss. | Mild cerebellar volume loss and focal gliosis in the right cerebllar hemiphere. | 10 | Intermittent generalized episodes of clonus and dystonic tremor | x |  |  |  |  |  |  |  |  | 0 |
| M | 0.52 | generalized | Prematurity | Abnormal basal ganglia signal. | Deep cerebellar/cerebellar nuclear signal abnormality | 2 | Generalized dystonia |  |  | x |  |  |  |  |  |  | 0 |
| F | 0.1 | generalized | HIE | Encephalomalacia of bilateral cerebral hemispheres, basal ganglia, and thalami. | Diffuse cerebellar volume loss | 0.25 | all extremities with opisthotonic postures |  |  |  |  |  | x |  |  |  | 7 |
| F | 1 | generalized | HIE | Encephalomalacia of cerebral hemispheres, basal ganglia, and thalami. | Cerebellar atrophy and gliosis. | 2.375 | Opisthotonic postuing with left version | x |  | x |  |  |  |  |  |  | 0 |
| F | 5 | focal | NMDA Encephalitis | Mild cerebral volume loss. | Mild cerebellar volume loss. | 5 | Provokable episodic dystonia |  |  |  |  |  | x |  |  |  | ? |
| M | 0.58 | generalized | Kernicterus | Basal ganglia and hippocampal volume loss and focal left thalamic encephalomalacia | Cerebellar volume loss |  | Not characterized |  |  |  |  |  |  |  |  | x | 1 |
| F | 14 | focal segmental | Anoxic Brain Injury | Cerebral volume loss and abnormal signal in basal ganglia and thalami. | Cerebellar atrophy | 14 | Rigid dystonic posturing |  |  |  |  |  | x |  |  |  |  |
| M | 1 | generalized | HIE | Cerebral encephalomalacia and abnormal basal ganglia and thalamic signal. | Cerebellar atrophy and signal change. | 1.25 | Opisthotonic posturing. |  |  |  | x |  |  |  |  |  | 1 |
| % M | 1 | generalized | Prematurity | Basal ganglia and thalamic volume loss | greater than right, abnormal signal in the dentate nuceli. | 18 | Dystonic lower extremity movements |  | x |  |  |  |  |  |  |  | 5 |
| & F | 1 | generalized | Prematurity | Cerebral volume loss. | Bilateral Cerebellar hemiphere encephalomalacia and severe atrophy. Pontine atrophy. | 18 | trunk present in episodes with spasticity |  |  |  |  |  |  |  |  |  | 9 |
| £ M | 0.5 | generalized | Perinataly Injury | Focal occipital and parietal encephalomalacia | Bilateral cerebellar encephalomalacia. | 7 | Intermittent episodes with preserved range of motion | x |  |  |  |  |  |  |  |  | 0 |
| M | 1.75 | generalized | In Utero (SOD) | Corpus callosal hypoplasia | Cerebellar atrophy | 1.75 | Increased axial and upper extremity tone |  |  |  |  |  |  | x |  |  | 3 |
| M | 13 | focal segmental | Traumatic Brain Injury | Basal ganglia encephalomalacia. | Cerebellar signal abnormalities | 13 | Generalized dystonia of upper greater than lower extremities |  |  |  |  |  |  |  | x |  | 1 |
| M | 16 | generalized | Infectious (West Nile virus) | Diffuse signal abnormalities of basal ganglia, thalamus, and cortex. | Atrophy of the left greater than right cerebellar hemiphere. | 18 | Upper extremities held in flexion |  |  |  |  |  |  |  |  |  | 1 |
| M | 10 | generalized | HIE | Cerebral, basal ganglia, and thalamic signal abnormalities | Cerebellar atrophy and signal change. | 16 | Rigid upper extremity flexion lower extremity extension | x |  | x |  |  |  |  |  |  | 3 |
| € F | 1.92 | focal segmental | NMDA Encephalitis | Normal imaging | Left focal cerebellar encephalomalacia | 1.92 | Opisthotonic posture with right version | x |  |  |  |  |  |  |  |  | 0 |
| F | 3 | generalized | Prematurity | Cerebral volume loss | Atrophic cerebellum with hypoplastic pons. | 10 | Opisthotonic extension with right version | x |  |  | x |  |  |  |  |  | 3 |
| M | 1 | generalized | HIE | Bilateral cerebral and deep gray encephalomalacia. | Cerebellar volume loss and gliosis with prominent cerebellar folia. | 1.17 | Variable dystonic movements observed | x |  | x |  |  |  |  |  |  | 1 |

### Photoactivation of the iCNN pathway induces dystonic crises reminiscent of the human condition in a mouse model of dystonia, but not in healthy mice

Although cerebellar nuclei neurons (*19*, *28*) have been implicated in driving the baseline symptoms of dystonia, it remains unclear whether cerebellar nuclei neurons contribute to dystonic crisis. As previous findings and our clinical results support a potential role for inhibitory cerebellar neurons, we hypothesized that iCNNs specifically instigate dystonic crisis. To test this, we turned to animal models of dystonia as a tractable means of determining the role of iCNNs in dystonic crisis. As previous studies showed increased activity of iCNNs in an animal model of dystonia during paroxysmal episodes of dystonic symptoms, we reasoned that optogenetic activation of iCNNs could induce the more severe dystonic crisis in an animal model of dystonia. To test this, we devised a mouse genetic model that could simultaneously also be manipulated with optogenetics. Since the development of dystonic crises almost exclusively occur in patients with a pre-existing diagnosis of dystonia (*5*, *29*), we sought to optogenetically manipulate the iCNN pathway in a mouse model with robust dystonic symptoms. The *Ptf1a^Cre^;Vglut2^fx/fx^* mouse (*30*, *31*) was chosen because of the cerebellar specificity of the manipulation that gives rise to dystonia, its *Ptf1a*-specific expression of Cre, and the severity of its dystonic symptoms (**Figure 2A**). As *Ptf1a* is expressed in iCNNs among other areas (*32–34*), it allowed cre-inducible expression within iCNNs to manipulate their activity and visualize their anatomy with genetic precision. In these mutant mice, vesicular glutamate transporter 2 (VGlut2) is genetically excised in *Ptf1a*-expressing neurons. As only inferior olive neurons, a major cerebellar input to cerebellar Purkinje cells, express Ptf1a and Vglut2 (*32*), this manipulation selectively silences neurotransmission from olivary climbing fibers to the cerebellum (**Figure 2A**), resulting in a robust dystonic phenotype described previously (*30*). These mutant mice exhibit dystonic symptoms of mild-to-moderate twisting postures in the limbs, trunk, head, and/or tail interrupted by random and intermittent periods of severe generalized dystonia in several body parts reminiscent of human dystonic crises. Our previous work has shown that the dystonic phenotype of *Ptf1a^Cre^;Vglut2^fx/fx^* mice is mediated by co-contractions of agonist-antagonist muscle pairs (*31*) and not seizures (*30*), making them an ideal model in which to investigate whether iCNN activity triggers dystonic crisis. Similar to human patients in which the same underlying genetic mutation may result in a spectrum of dystonic symptoms across patients (*35*, *36*), the severity of the underlying dystonic symptoms in the *Ptf1a^Cre^;Vglut2^fx/fx^*mouse can vary between animals. To selectively manipulate the iCNN pathway in these mice using optogenetics, we crossed the *Ptf1a^Cre^;Vglut2^fx/fx^*mice with *ROSA^lsl-Chr2/EYFP^* mice to generate *Ptf1a^Cre^;Vglut2^fx/fx^;ROSA^lsl-Chr2/EYFP^* mice (**Figure 2B**). To confirm that iCNNs selectively expressed the ChR2 construct in the *Ptf1a^Cre^;Vglut2^fx/fx^;ROSA^lsl-Chr2/EYFP^* mice, we performed *in situ* hybridization assays for the YFP tag in the construct, *Vgat* (inhibitory marker), and *Vglut2* (excitatory marker). Robust co-labeling of YFP and *Vgat*, but not *Vglut2*, was observed in the mutant deletion mice (**Figure 2C** and **Supplemental Figure 1**, n=3 animals). To selectively manipulate the iCNN pathway and avoid activation of Purkinje cell terminals (which also express *Ptf1a*) in the cerebellar nuclei, we targeted iCNN axons by bilaterally implanting optical fibers over the superior cerebellar peduncle, the white matter tract through which extracerebellar cerebellar nuclei neuron projections pass (*17*) (**Figure 2D** and **Supplemental Figure 2**). The selective manipulation of cerebellar nuclei neurons by targeting the superior cerebellar peduncle has been similarly employed to create an animal lesion model of dystonic crisis (*18*) and to therapeutically treat refractory dystonia in patients (*37*, *38*). The behavior of experimental mice in an open field was video recorded and data from two minutes before and during stimulation were analyzed (470 nm light; 50 Hz square pulses; ≥ 1mW at fiber tip). To detect changes in the incidence of dystonic crises, we measured the percentage of time that a mouse spent in a dystonic crisis during each period (before vs during stimulation). Throughout this study, we define a dystonic crisis in mice as observable dystonic limb contractions, limb hyperextensions and/or overt postures in at least two body parts in addition to severely limited ambulation for at least 3 seconds, a sufficient time to confidently determine that ambulation was restricted (rather than due to voluntary resting) and a result of abnormal muscle activity (**<u>Supplementary Movie 2</u>**).

**Figure 2:**
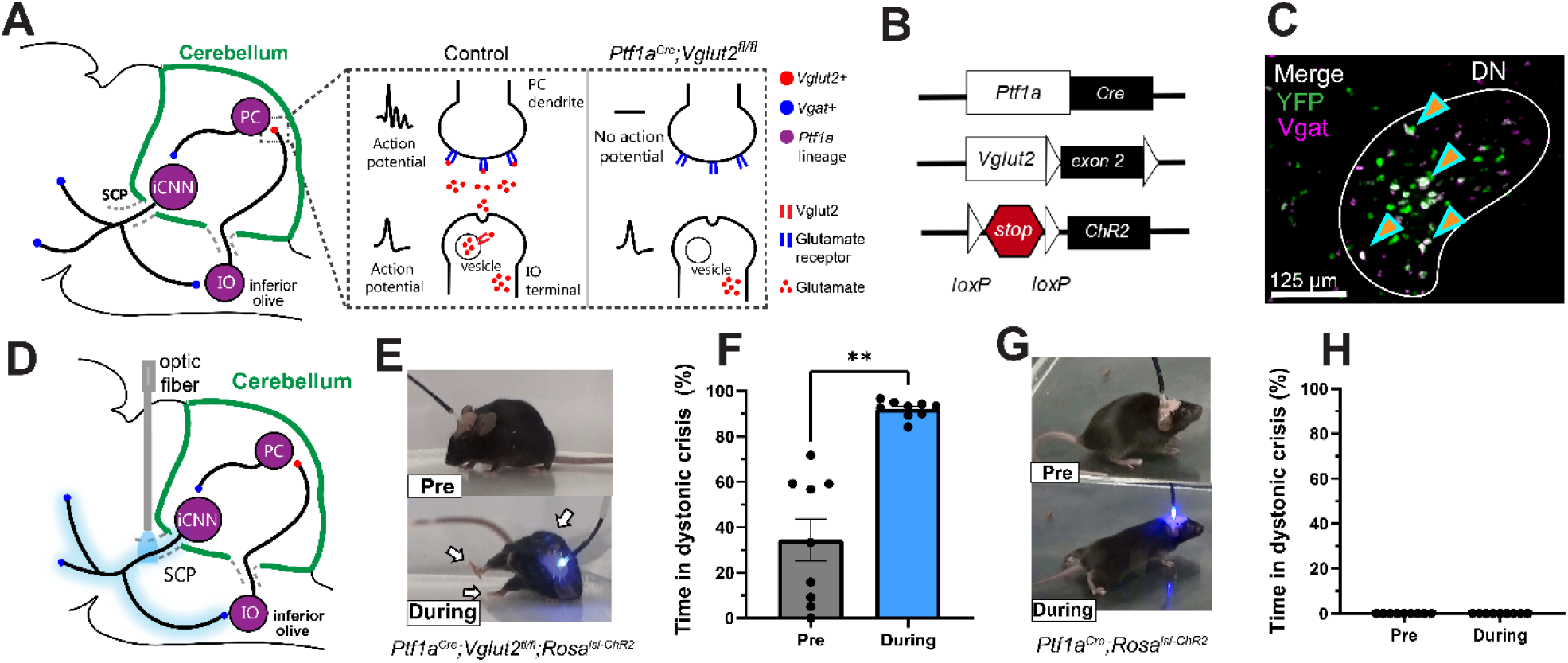
Photoactivation of the inhibitory cerebellar nuclei neuron (iCNN) pathway induces dystonic crises reminiscent of the human condition in *Ptf1a^Cre^;Vglut2^fx/fx^;ROSA^lsl-Chr2/EYFP^*mice with existing dystonia, but not in otherwise healthy mice. **(A)** Schematic showing the genetic manipulation in *Ptf1a^Cre^;Vglut2^fx/fx^* mice that selectively silenced olivocerebellar synapses, which causes dystonia. (B) Using the *Ptf1a^Cre^* driver, exon 2 of *Vglut2* was selectively removed and *Vglut2* deleted from the inferior olive. Concurrently, the loxP sites surrounding a stop signal preceding ChR2-YFP were removed, allowing expression of ChR2-YFP in Ptf1a-expressing neurons including inferior olivary neurons, Purkinje cells, and iCNNs. (C) *In situ* image showing co-labeling of *YFP* and *Vgat* in the cerebellar nuclei. (D) Schematic showing the selective photoactivation iCNNs through their projections in the superior cerebellar peduncle. (E) Photoactivation of the iCNN pathway induced a dystonic crisis on-demand. (F) In *Ptf1a^Cre^;Vglut2^fx/fx^;ROSA^lsl-Chr2/EYFP^*mice, photoactivation of the iCNN pathway significantly increased the percentage of time spent in dystonic crisis when comparing before and during stimulation (n=9 animals; p=0.0039, Wilcoxon matched-pairs signed rank test, two-tailed). (G) Photoactivation of the iCNN pathway in healthy *Ptf1a^Cre^;ROSA^lsl-Chr2/EYFP^*mice resulted in abnormal movements, but not dystonic crises. (H) Photoactivation of the iCNN pathway did not influence the percentage of time spent in dystonic crisis when comparing before and during stimulation (n=9 animals). Error bars are defined as standard error of the mean. SCP, superior cerebellar peduncle; PC, Purkinje cell; iCNN, inhibitory cerebellar nuclei neuron; IO, inferior olive; DN, dentate cerebellar nucleus.

Photoactivation of the iCNN pathway induced severe dystonic crises on-demand which was clearly observable in videos (**Figure 2E** and **<u>Supplementary Movie 2</u>**; i.e. extended limbs, rigidity throughout the body, stiff tail, splayed digits, toppling over, severely impaired ambulation). The photoinduced dystonic crises were similar to the most severe spontaneous periods of dystonia observed in these mutant mice and reminiscent of the acute, severe episodes of dystonia observed in another murine model of dystonic crisis (*18*). The photoactivation period was marked by a significant increase in the percentage of time spent in a dystonic crisis during stimulation (92%) compared to before stimulation (34%), supporting that a dystonic crisis was induced (**Figure 2F**, n=9 animals; p=0.0039, Wilcoxon matched-pairs signed rank test, two-tailed). The photoinduced dystonic crisis occurred within 9 seconds of stimulation onset (**Supplemental Figure 3A**; n=9 animals), persisted throughout the duration of the light stimulation, and did not end until 38 seconds after the stimulation ended (**Supplemental Figure 3B**; n=9 animals).

Although dystonic crisis occurs in patients with pre-existing dystonia in most cases, it is possible for otherwise heathy patients to develop a dystonic crisis (*5*, *29*). Likewise, it may be possible that the iCNN pathway may broadly generate general dystonic movements rather than specifically trigger dystonic crisis in the context of a pre-existing dystonic condition. To test this, we repeated this optogenetic experiment in otherwise heathy mice with an opsin (*Ptf1a^Cre^;ROSA^lsl-Chr2/EYFP^*). Photoactivation of the iCNN pathway in healthy mice (*Ptf1a^Cre^;ROSA^lsl-Chr2/EYFP^*) did not result in dystonic crises (**Figure 2G** and **<u>Supplementary Movie 3</u>**) as the percentage of time in crisis remained 0 before and during the stimulation (**Figure 2H**; n=9 animals). However, iCNN photoactivation did lead to abnormal movement and short, intermittent, mild dystonic movements. Almost instantaneously with the onset of stimulation, the mouse was observed to be more tense, lowered its body to the ground as if crawling, widened its hind legs, dragged its trunk along the floor while walking, and appeared to “waddle” with slow and clunky locomotion forward and wide steps (**Figure 2G** and **<u>Supplementary Movie 3</u>**). These data suggest that the pre-existing dystonic state of *Ptf1a^Cre^;Vglut2^fx/fx^;ROSA^lsl-Chr2/EYFP^*mice sensitizes the iCNN pathway to trigger dystonic crises. As dystonic crisis predominantly arises in patients with a pre-existing diagnosis of dystonia, these results further support a role for the iCNN pathway in the origin of dystonic crises.

### Photoinhibition of the iCNN pathway alleviates spontaneous dystonic crises in a mouse model of dystonia

As photoactivation of the iCNN pathway induced dystonic crisis, we reasoned that photoinhibition of the iCNN pathway would alleviate spontaneous dystonic crises in *Ptf1a^Cre^;Vglut2^fx/fx^* mice. To test this, we generated *Ptf1a^Cre^;Vglut2^fx/fx^;ROSA^lsl-Arch^*mice expressing the inhibitory opsin, archaerhodopsin, and targeted photostimulation to the iCNN pathway as previously described (**Figure 3A**). As we were interested in detecting a reduction in the time spent in dystonic crisis and the severity of dystonia varies across individual mice, we selected *Ptf1a^Cre^;Vglut2^fx/fx^;ROSA^lsl-Arch^*mice that had severe dystonic crises frequently. To address the presence of possible lags in the onset of therapeutic effects of photoinhibition, as previously observed in treatments of dystonic patients (*39–42*) and animal models (*30*), we repeated the experiment over four consecutive days. To establish a baseline for how much time the mice spend in dystonic crisis over a 35-minute session without optogenetic manipulation, we included a sham-stimulation day on Day 0, in which the mice were connected to the optic fibers throughout the 35-minute session as on subsequent days, but no light stimulation was administered (**Figure 3B**). The sham-stimulation day on Day 0 was an important control to distinguish whether changes in behavior during the 35-minute session may be instigated by some other confounding variable that may be independent of optogenetic manipulation, such as time, stress, environmental familiarity, handling, or restraint by the optic fibers coupled to the head of an experimental mouse. To test whether photostimulation influenced the percentage of time spent in dystonic crisis, video was recorded for five minutes before and five minutes during stimulation (561 nm light; continuous light; ≥ 1mW at fiber tip). To increase confidence that any behavioral change was a result of optogenetic manipulation, we repeated the stimulation paradigm three times on each day, resulting in a 35-minute session for each day (**Figure 3B**). For each day, we averaged the percentage of time spent in crisis before and during the stimulation periods and then compared the resulting values for statistical significance.

**Figure 3:**
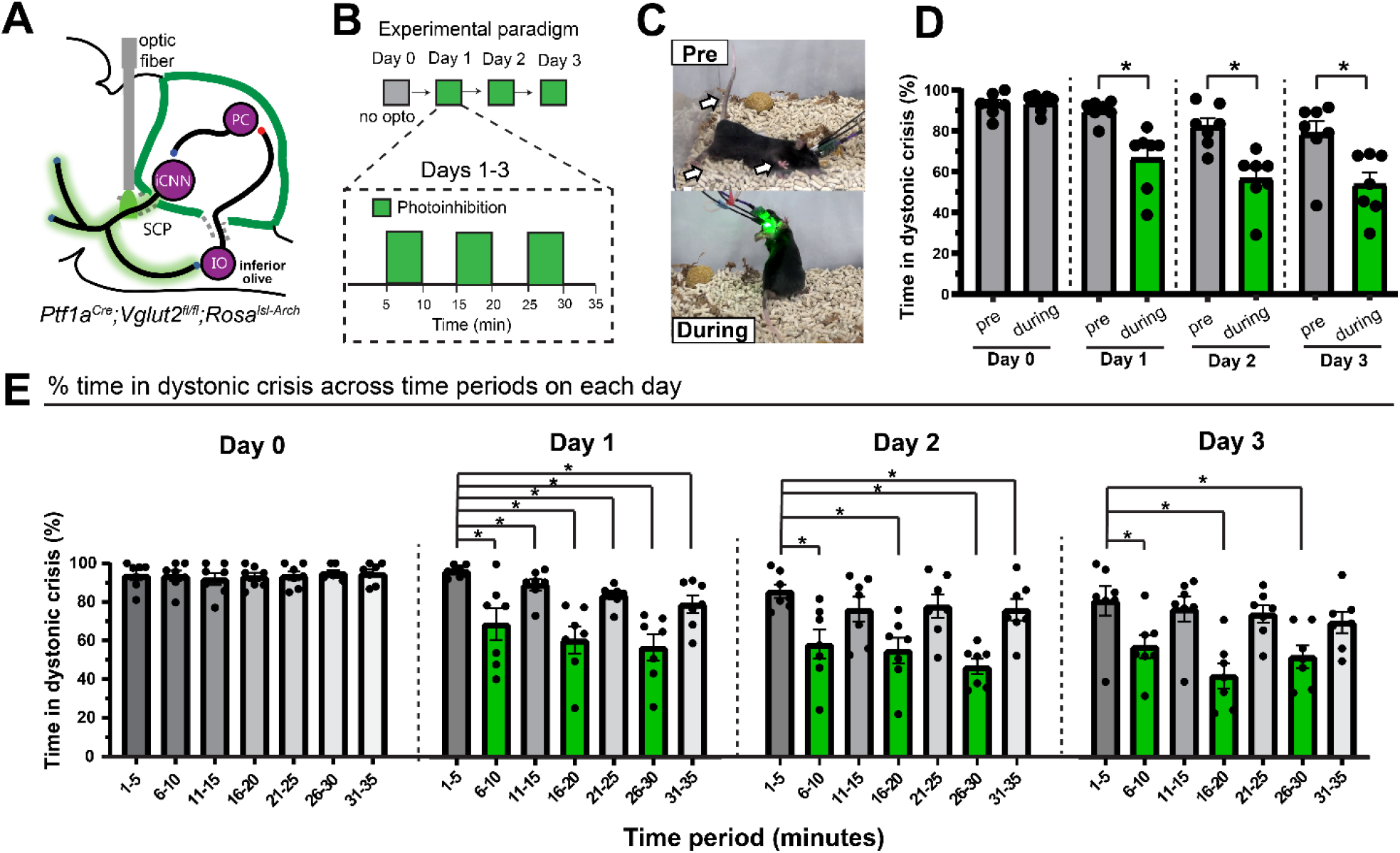
Photoinhibition of the inhibitory cerebellar nuclei neuron (iCNN) pathway alleviates the spontaneous dystonic crises in *Ptf1a^Cre^;Vglut2^fx/fx^;ROSA^lsl-Arch/EYFP^*mice with existing dystonia. **(A)** Schematic showing the selective photoinhibition of the iCNN pathway by implanting optical fibers into the superior cerebellar peduncle of *Ptf1a^Cre^;Vglut2^fx/fx^;ROSA^lsl-Arch^*mice. (B) Schematic showing the four-day experimental paradigm. (C) Photoinhibition of the iCNN pathway alleviated dystonic crises in *Ptf1a^Cre^;Vglut2^fx/fx^;ROSA^lsl-Arch^* mice as shown in (D) by a significant decrease in the percentage of time spent in crisis when comparing pre versus during photostimulation on Day 1 (p=0.0156), Day 2 (p=0.0156), and Day 3 (p=0.0156) (E) Photoinhibition on Day 1 reduced dystonic crises across all time periods (for 1-5 vs 6-20, p=0.0312; for 1-5 vs 11-15, p=0.0156; for 1-5 vs 16-20, p=0.0156; for 1-5 vs 21-25, p=0.0156; for 1-5 vs 26-30, p=0.0156; for 1-5 vs 31-35, p=0.0156). Photoinhibition on Day 2 alleviated crises across several time periods (for 1-5 vs 6-20, p=0.0156; for 1-5 vs 11-15, p=0.2345; for 1-5 vs 16-20, p=0.0156; for 1-5 vs 21-25, p=0.2969; for 1-5 vs 26-30, p=0.0156; for 1-5 vs 31-35, p=0.0469). Photoinhibition on Day 3 reduced crises across several time periods (for 1-5 vs 6-20, p=0.0156; for 1-5 vs 11-15, p=0.2188; for 1-5 vs 16-20, p=0.0156; for 1-5 vs 21-25, p=0.1562; for 1-5 vs 26-30, p=0.0156; for 1-5 vs 31-35, p=0.0781). For all statistics: n=7 animals, Wilcoxon matched-pairs signed rank test, two-tailed, error bars are defined as standard error of the mean.

To test whether photoinhibition of the iCNN pathway led to a same-day reduction in dystonic crises, we first established a baseline by performing the experiment without any photostimulation on Day 0 - the sham-stimulation day (**Figure 3B**). As expected on Day 0 when no photostimulation was given, we did not find a significant change in the percentage of time spent in dystonic crisis when comparing periods corresponding to before and during stimulation (93% vs 94%; p=0.6875) in severely dystonic *Ptf1a^Cre^;Vglut2^fx/fx^;ROSA^lsl-Arch^* mice (**Figure 3D**, n=7 animals, Wilcoxon matched-pairs signed rank test, two-tailed). Remarkably, same-day photoinhibition of the iCNN pathway alleviated spontaneous dystonic crises and rescued mobility in these mice (**Figure 3C** and **<u>Supplementary Movie 4</u>**) as shown by a significant decrease in the percentage of time spent in dystonic crisis when comparing before versus during photostimulation on Day 1 (90% vs 66%; p=0.0156), Day 2 (82% vs 57%; p=0.0156), and Day 3 (79% vs 54%; p=0.0156)(**Figure 3D**; n=7 animals, Wilcoxon matched-pairs signed rank test, two-tailed).

As our previous analysis for each day averaged the effects of three stimulations over 35 minutes, we wondered whether the observed therapeutic effects were triggered immediately by the first stimulation or whether they developed progressively over the course of the three stimulations of each day. To test this, the percentage of time in dystonic crisis across five-minute periods on a given day was compared (**Figure 3E**). On Day 0 (sham-stimulation), when comparing the first five minute period (1–5) to each subsequent five-minute time period (6–10, 11–15, 16–20, 21–25, 26–30, 31–35), there was no significant change (**Figure 3E**, n=7 animals; for 1-5 vs 6-20, p=0.7500; for 1-5 vs 11-15, p=0.0625; for 1-5 vs 16-20, p=0.9219; for 1-5 vs 21-25, p=0.8750; for 1-5 vs26-30, p=0.6719; for 1-5 vs 31-35, p=0.8438; Wilcoxon matched-pairs signed rank test, two tailed), suggesting that spontaneous dystonic crises do not change across the 35-minute session. This suggests that possible confounding variables, such as environmental familiarity or absence of handling, likely do not influence the expression of dystonic crises in the experimental mice. When making the same comparison on Day 1, we found a significant reduction in dystonic crises across all time periods (**Figure 3E**, n=7 animals; for 1-5 vs 6-20, p=0.0312; for 1-5 vs 11-15, p=0.0156; for 1-5 vs 16-20, p=0.0156; for 1-5 vs 21-25, p=0.0156; for 1-5 vs 26-30, p=0.0156; for 1-5 vs 31-35, p=0.0156; Wilcoxon matched-pairs signed rank test, two tailed). On Day 2, we found a significant reduction in dystonic crisis across several time periods (**Figure 3E**, n=7 animals; for 1-5 vs 6-20, p=0.0156; for 1-5 vs 11-15, p=0.2345; for 1-5 vs 16-20, p=0.0156; for 1-5 vs 21-25, p=0.2969; for 1-5 vs 26-30, p=0.0156; for 1-5 vs 31-35, p=0.0469; Wilcoxon matched-pairs signed rank test, two tailed). Day 3 showed a similar significant reduction during stimulation periods (**Figure 3E**, n=7 animals; for 1-5 vs 6-20, p=0.0156; for 1-5 vs 11-15, p=0.2188; for 1-5 vs 16-20, p=0.0156; for 1-5 vs 21-25, p=0.1562; for 1-5 vs 26-30, p=0.0156; for 1-5 vs 31-35, p=0.0781; Wilcoxon matched-pairs signed rank test, two tailed). As the significant reductions in the time spent in dystonic crisis on Day 2 and Day 3 predominantly occur during stimulation periods and not non-stimulation periods, they support the idea that the alleviation of dystonic crises is a consequence of our optogenetic manipulation rather than other confounding variables like exhaustion or familiarity with the environment. These results further suggest that photoinhibition of the iCNN pathway has short-term, same-day, cumulative therapeutic effects on dystonic crisis.

To explore whether any long-term, multiday, cumulative therapeutic effects may have occurred across days, the percentage of time in dystonic crisis was compared across the pre stimulation periods and during stimulation periods across days (**Supplemental Figure 4**). For the pre stimulation periods when comparing Day 1 (the first day of stimulation) against Day 2 and Day 3, we report a trend toward a reduction in dystonic crises, but it is not statistically significant (**Supplemental Figure 4A,** n=7 animals; for Day 1 vs Day 2, n=0.0625; for Day 1 vs Day 3, n=0.0781; Wilcoxon matched-pairs signed rank test, two tailed). For the during stimulation periods, we found a significant reduction in dystonic crises when comparing Day 1 to Day 2 and Day 3 (**Supplemental Figure 4B,** n=7 animals; for Day 1 vs Day 2, p=0.0312; for Day 1 vs Day 3, p=0.0156; Wilcoxon matched-pairs signed rank test, two tailed). These results suggest that the photoinhibition of the iCNN pathway also has long-term, multiday, cumulative therapeutic effects on dystonic crisis.

Together, these results suggest that photoinhibition of iCNN activity immediately alleviates spontaneous dystonic crises in mice with severe dystonia, and that these beneficial therapeutic effects accumulate over time.

### iCNNs send direct projections to the centrolateral nucleus of the thalamus (CL)

As dystonia is thought to result from aberrant interactions across structures in the motor network (*43*), we reasoned that the iCNN pathway interacts with other structures in the motor network to control the onset of a dystonic crisis. Emerging evidence in patients (*35*, *43–46*) and animal models (*30*, *47*, *48*) support a role for the cerebello-thalamo-basal ganglia pathway in several forms of dystonia, and anatomical studies in non-human primates and animal models have revealed that the cerebellar nuclei interact with the basal ganglia through the CL (*49*, *50*). However, it is thought that these cerebello-thalamic projections predominantly originate from excitatory, not inhibitory, cerebellar nuclei neurons (*20*, *50*, *51*). Moreover, it remains unknown whether the cerebello-thalamo-basal ganglia pathway contributes to dystonic crisis. To investigate whether iCNNs form a part of the cerebello-thalamo-basal ganglia pathway, we hypothesized that there may be a previously unknown monosynaptic projection from iCNNs to the CL. In *Ptf1a^Cre^;Vglut2^fx/fx^;ROSA^lsl-Chr2/EYFP^* mice, we identified axons labeled with ChR2-EYFP in the CL (**Figure 4A**), suggesting that there may indeed be monosynaptic iCNN projections. However, as these projections could originate from any *Ptf1a*-expressing region (including the hypothalamus, retina, spinal cord, cerebellum, brainstem, or forebrain), we designed an experiment to determine whether iCNNs have a robust monosynaptic projection to the CL using a combination of intersectional genetics and injected neural tracers. To genetically label only the cell bodies of *Ptf1a*-expressing neurons like iCNNs, we crossed *Ptf1a^Cre^;Vglut2^fx/fx^* with a cre-dependent Sun1 (a small nuclear envelope protein) reporter line to generate *Ptf1a^Cre^;Vglut2^fx/fx^;ROSA^lsl-Sun1-GFP^* mice (**Figure 4B**). To retrogradely label the cell bodies of iCNNs that may project to the CL, we bilaterally injected BDA 3K (a predominantly retrograde neural tracer (*52*, *53*)) into the CL (**Figure 4C-D**). We found co-labeling of DAPI, BDA 3K, and Sun1-GFP in the cerebellar nuclei (**Figure 4E, inset i; orange ticks**), suggesting that iCNNs monosynaptically project to the CL. We found co-labeling of these three markers across all three major cerebellar nuclei comprising the fastigial, interposed, and dentate cerebellar nuclei (**Supplemental Figure 5**). These data identify a specific cerebello-thalamic anatomical pathway through which iCNNs could control and/or contribute to the onset of dystonic crisis.

**Figure 4:**
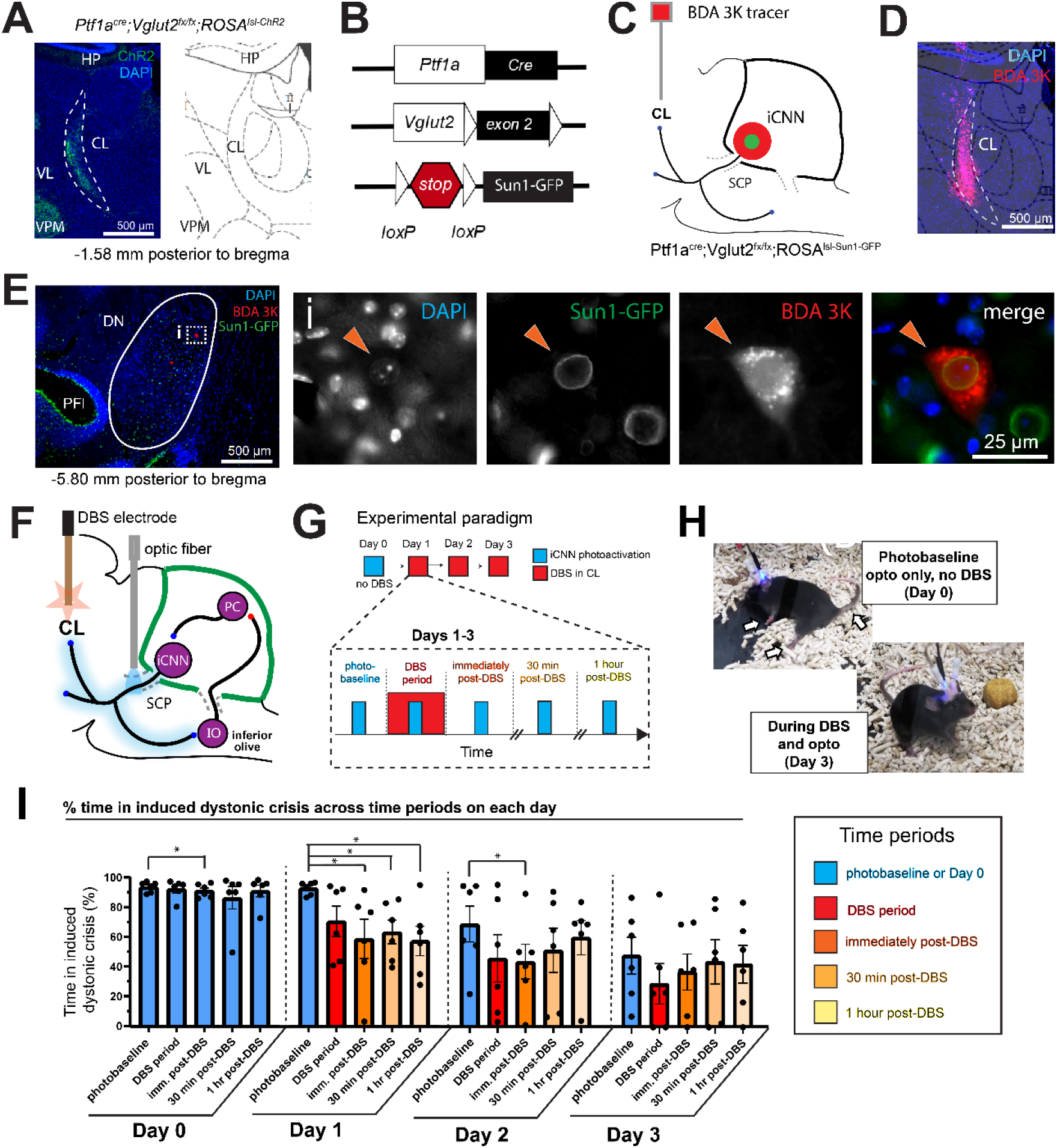
Inhibitory cerebellar nuclei neurons (iCNNs) interact with the centrolateral nucleus of the thalamus (CL) during dystonic crisis. **(A)** Immunohistochemical staining in *Ptf1a^Cre^;Vglut2^fx/fx^;ROSA^lsl-Chr2/EYFP^*mice revealed ChR2-YFP-labeled projections in the CL. **(B)** Schematic of genetics in *Ptf1a^Cre^;Vglut2^fx/fx^;ROSA^lsl-Sun1-GFP^* mice that allowed localized GFP expression in the somas of Ptf1a-expressing neurons. (C) Experimental setup to identify iCNNs projecting to the CL. Retrograde BDA 3K tracer was injected into CL of *Ptf1a^Cre^;Vglut2^fx/fx^;ROSA^lsl-Sun1-GFP^*mice as in (D), and iCNNs were co-labeled with BDA 3K and GFP as shown in (E, inset i) (n=4 mice). (F) Schematic showing the selective photoactivation of iCNN projections in the SCP and DBS of the CL. (G) Four-day experimental setup showing the optogenetic and DBS paradigms during the five distinct time periods for each day. (H) Representative images showing how DBS alleviated dystonic crisis induced by photoactivation of the iCNN pathway. (I) Dystonic crises induced by photoactivation of the iCNN pathway did not change greatly across time periods on Day 0 though a statistical difference was detected when comparing the photobaseline to immediately post-DBS (p=0.0312). Induced crises were substantially reduced on Day 1 when comparing the photobaseline to immediately post-DBS (p=0.0312), 30 minutes post-DBS (p=0.0312), and 1 hour post-DBS (p=0.0312) and on Day 2 when comparing the photobaseline to immediately post-DBS (p=0.0312). For all statistics: n=6 animals, Wilcoxon matched-pairs signed rank test, two-tailed, error bars are defined as standard error of the mean. CL, centrolateral nucleus of the thalamus; HP, hippocampus; VL, ventrolateral thalamus; VPM, ventroposterior thalamus; Pfl, paraflocculus; DN, dentate cerebellar nucleus. SCP, superior cerebellar peduncle; DBS, deep brain stimulation; iCNN, inhibitory cerebellar nuclei neuron.

### Photoactivation of iCNN-CL projections modestly increases time spent in dystonic crises

As our data showed that iCNN photoactivation induces dystonic crises and that iCNNs project to the CL, we sought to test whether iCNNs and the CL interact to control the onset of dystonic crisis and reflect a possible therapeutic node. To test whether selective activation of iCNN-CL projections may be sufficient to induce dystonic crises as observed when targeting the entire iCNN pathway through the superior cerebellar peduncle, we photoactivated iCNN-CL terminals in *Ptf1a^Cre^;Vglut2^fx/fx^;ROSA^lsl-Chr2/EYFP^*mice (**Supplemental Figure 6A**). To better control our ability of optogenetically inducing dystonic crises even when the severity of dystonia varies across individual mice, we selected *Ptf1a^Cre^;Vglut2^fx/fx^;ROSA^lsl-ChR2/EYFP^*mice that had infrequent dystonic crises for this set of experiments. In this way, we would avoid a potential ceiling effect (e.g., where mice are so frequently in dystonic crisis that we cannot detect a worsening effect) and be able to better observe whether photostimulation may lead to a dystonic crisis. As in previous experiments, we conducted the experiment over four days (**Supplemental Figure 6B**) and included a sham-stimulation day on Day 0, in which the mice were connected to the optic fibers throughout the 35-minute session as on subsequent days, but no light stimulation was administered (**Supplemental Figure 6B**). As explained previously, the sham-stimulation day on Day 0 was an important control to distinguish whether changes in behavior during the 35-minute session may be instigated by some other confounding variable that may be independent of optogenetic manipulation. To test whether photostimulation influenced the percentage of time spent in dystonic crisis, video was recorded for five minutes before and five minutes during stimulation (480 nm light; 50 Hz square pulses; ≥ 1mW at fiber tip). To increase confidence that any behavioral change was a result of optogenetic manipulation, we repeated the stimulation paradigm three times on each day, resulting in a 35-minute session for each day (**Supplemental Figure 6B**). For each day, the percentage of time spent in dystonic crisis before and during the stimulation periods were averaged and statistically compared.

As expected on Day 0 when no photostimulation was given, there was no significant change in the percentage of time spent in dystonic crisis when comparing periods corresponding to before and during stimulation (13% vs 13%; p=0.8125 in *Ptf1a^Cre^;Vglut2^fx/fx^;ROSA^lsl-ChR2/EYFP^*mice (**Supplemental Figure 6C,** n=7 animals, Wilcoxon matched-pairs signed rank test, two-tailed). Photoactivation of the CNN-CL terminals modestly increased the time spent in dystonic crisis in these mice as shown by a modest but statistically significant increase in the percentage of time spent in dystonic crisis when comparing before versus during photostimulation on Day 1 (13% vs 21%; p=0.0156), Day 2 (14% vs 25%; p=0.0156), and Day 3 (14% vs 21%; p=0.0156). (**Supplemental Figure 6C**; n=7 animals, Wilcoxon matched-pairs signed rank test, two-tailed). These results suggest that the iCNN-CL projection can contribute to the incidence of dystonic crises, but not to the same extent as that induced by the photoactivation of widespread extracerebellar projections in the iCNN pathway through the superior cerebellar peduncle. This may implicate a synergistic role for cerebellar function and activity from additional downstream regions in line with the motor network theory of dystonia.

### DBS of the CL alleviates dystonic crises induced by iCNN photoactivation

Given that activating iCNN axon terminals in the CL modestly included dystonic crisis, we next wondered whether CL activity is required to induce the dystonic crises triggered by stimulating the broader iCNN pathway. This importance of CL activity in dystonia has been highlighted by previous studies showing that the disruption of CL activity with pharmacological agents, electrolytic lesions, or DBS can alleviate dystonic symptoms in animal models of dystonia (*18*, *30*, *47*, *50*) including the *Ptf1a^Cre^;Vglut2^fx/fx^*model. Thus, we reasoned that if the iCNN pathway, specifically, induces dystonic crises through the CL, then DBS of the CL will alleviate the dystonic crises induced by iCNN photoactivation.

To test this in the *Ptf1a^Cre^;Vglut2^fx/fx^;ROSA^lsl-Chr2/EYFP^*mice, we unilaterally implanted an optical fiber over the superior cerebellar peduncle and bilaterally implanted bipolar DBS electrodes into the CL (**Figure 4F** and **Supplemental Figure 7**). To account for possible delays in the therapeutic effect of DBS as previously observed in patients and animal models, we repeated the experiment over four consecutive days. Each day consisted of an 86-minute session divided into five distinct periods: photobaseline, DBS period, immediately post-DBS, 30 minutes post-DBS, and 1 hour post-DBS (**Figure 4G**). During each distinct period, a 2-minute stimulation of blue light (470 nm light; 50 Hz square pulses; ≥ 1mW at fiber tip) was administered followed by 2 minutes of no light stimulation. This paradigm was repeated three times for each distinct period. The percentage of time spent in a dystonic crisis was quantified for each of the three light pulses, and the average of these values was analyzed. To establish a baseline for how much time the mice spend in dystonic crises induced by iCNN photoactivation, we included a sham-DBS day on Day 0, in which the mice were connected to the stimulus generator throughout the 86-minute session as on subsequent days, but no DBS was administered (**Figure 4G**). As explained previously, the sham-stimulation day on Day 0 was an important control to distinguish whether changes in behavior during the 86-minute session may be instigated by some other confounding variable that may be independent of DBS. We administered DBS on Days 1-3. We only performed DBS in the DBS period for 10 minutes (130 Hz, 60µs, ±30µA), starting with the onset of the first light pulse and ending with the offset of the third light pulse. To use consistent language with days on which DBS was administered, we refer to the time periods on Day 0 as “Photobaseline,” “DBS period,” “Immediately post-DBS,” “30 minutes post-DBS,” and “1 hour post-DBS,” even though no DBS was administered on this day.

First, we analyzed whether CL-DBS led to same-day changes in the time in induced dystonic crisis across the five time periods (**Figure 4I**). During the sham-DBS stimulations on Day 0, the percentage of time in induced crisis across the five time periods remained mostly the same (between 83-94%), though there was a statistically significant reduction during the time period which would correspond to immediately post-DBS compared to the photobaseline (91% vs 94%;p=0.0312) (**Figure 4I**; n=6 mice, Wilcoxon matched-pairs signed rank test, two tails). This suggested that iCNN photoactivation reliably and consistently induced dystonic crises throughout the 86-minute session, and that little habituation occurs during repeated photostimulation of iCNNs with sham DBS on a single day. On Day 1, when comparing between the photobaseline and all subsequent time periods after DBS was administered, we found a marked reduction in the time spent in induced crisis when comparing the same-day photobaseline to during DBS (93% vs 70%; p=0.0625), immediately post-DBS (93% vs 58%; p=0.0312), 30 minutes post-DBS (93% vs 63%; p=0.0312), and 1 hour post-DBS (93% vs 57%; p=0.0312)(**<u>Supplementary Movie 5</u>** and **Figure 4I**, n=6 animals, Wilcoxon matched-pairs signed rank test, two tails). This progression suggested a short-term, delayed therapeutic effect of CL-DBS on iCNN-mediated dystonic crises. On Day 2, while the same general trend in reduced time in induced dystonic crisis following CL-DBS was observed, there was only a statistically significant reduction from the same-day baseline to immediately post-DBS (69% vs 44%; p=0.0312)(**Figure 4I**, n=6 animals, Wilcoxon matched-pairs signed rank test, two tails). On Day 3, we did not find statistically significant changes though the same general trend in reduced crisis time following DBS can be seen (**Figure 4I**). Importantly, a trending decrease in dystonic crises across days can be appreciated (**Figure 4H-I**), especially in the photobaseline periods, which may explain the lack of statistically significant comparisons on Days 2-3. These data suggest that CL-DBS causes long-term, multi-day, therapeutic changes that alleviate dystonic crises induced by iCNN photoactivation.

To further explore whether CL-DBS led to long-term, multi-day therapeutic changes in the induced dystonic crisis time, we compared the time in induced dystonic crisis for each period across Days 0-3 (**Supplemental Figure 8**). When comparing the photobaseline period across days (**Supplemental Figure 8A**), we observed a clear reduction in the time in induced crisis on Day 2 (69%) and Day 3 (48%) of CL-DBS compared to Day 0 (94%) of sham-DBS and Day 1 (93%) of CL-DBS. This corresponded to the days after CL-DBS had been administered at least once, suggesting that CL-DBS likely led to this long-term therapeutic change in dystonic crisis induction, rather than other confounding variables like habituation to photostimulation or familiarity with the environment. However, we only detected a statistically significant reduction during the photobaseline period when comparing Day 0 of sham-DBS with Day 3 of CL-DBS (**Supplemental Figure 8A**, n=6 animals, p=0.0312, Wilcoxon matched pairs signed rank test, two tails). For the DBS period, when comparing against the time in crisis on Day 0 (92%) of sham-DBS, we observed a trending but non-significant reduction on Day 1 (92% vs 70%; p=0.1562) of CL-DBS and a statistically significant reduction on Day 2 (92% vs 46%; p=0.0312) and Day 3 (92% vs 29%; p=0.0312) of CL-DBS (**<u>Supplementary Movie 5</u>** and **Supplemental Figure 8A**, n=6 animals, Wilcoxon matched-pairs signed rank test, two tails). For immediately post-DBS, the induced dystonic crisis time on Day 0 of sham-DBS significantly decreased on Day 1 (91% vs 58%; p=0.0312), Day 2 (91% vs 44%; p=0.0312), and Day 3 (91% vs 37%; p=0.0312) of CL-DBS (**Supplemental Figure 8A**, n=6 animals, Wilcoxon matched-pairs signed rank test, two tails). For 30 minutes post-DBS, the induced crisis time on Day 0 of sham-DBS decreased on Day 1 (86% vs 63%; p=0.0312), Day 2 (86% vs 51%; p=0.01562), and Day 3 (86% vs 44%; p=0.0312) of CL-DBS, though it only reached statistical significance for Days 1 and 3 of CL-DBS (**Supplemental Figure 8A**, n=6 animals, Wilcoxon matched-pairs signed rank test, two tails). For 1 hour post-DBS, the induced crisis time on Day 0 of sham-DBS decreased significantly on Day 1 (92% vs 61%; p-0.0156), Day 2 (92% vs 64%; p-0.0156), and Day 3 (93% vs 49%; p=0.0156) of CL-DBS (**Supplemental Figure 8A**, n=6 animals, Wilcoxon matched-pairs signed rank test, two tails). Together, these results suggest that the iCNN pathway and CL interact to control the onset of dystonic crises, and that CL-DBS may provide prolonged and cumulative therapeutic effects for alleviating dystonic crises.

## Discussion

The translational relevance of our study is suggested by the retrospective clinical cohort where more patients admitted for dystonic crisis (e.g., status dystonicus) presented with cerebellar lesions rather than supratentorial lesions and responded favorably to inhibitory drugs to alleviate dystonic crisis. Future clinical studies with larger cohorts could expand on these findings by controlling for potential confounding variables such as dystonia etiology and comorbidities. Additionally, it would be interesting to compare outcomes for patients with cerebellar-only versus cerebellar plus supratentorial lesions, and to analyze differences between patients with lesions in different cerebellar subregions. Although the current clinical results are correlative in nature, they led us to hypothesize that iCNNs may instigate dystonic crisis, which we tested in an animal model.

As previously described, dystonic crisis is a life-threatening medical emergency. Such intense dystonic contractions can lead to muscle breakdown or impaired breathing (*6*, *54*, *55*). Although our experiments were not designed to quantify these severe aspects of human dystonic crisis in our mouse model, our results show that photoactivating iCNNs can significantly increase the severity of dystonia, pushing mice from intermittent dystonic postures to constant dystonic immobility in a manner that closely models the emergence of dystonic crisis in patients as consistently described in the literature. Furthermore, our results show that inhibiting iCNNs has the opposite effect and reduces the frequency of severe dystonia. Together, these findings strongly indicate that iCNNs may be key to the development and treatment of dystonic crises. Additional studies specifically evaluating the role of iCNNs in dystonic crisis in humans will be critical in translating our findings into clinical practice.

Little is known about iCNNs, and their functional contribution has been historically limited to motor learning and coordination (*56–60*). However, recent studies have increasingly supported a more prominent role for this subtype of cerebellar nuclei neurons, showing that they have distinct anatomical, electrophysiological, and protein expression properties (*20*, *21*, *56*) and that they may be involved in non-motor processes (*61*). For example, Judd et al. 2021 found that iCNNs restricted to the anterior interposed cerebellar nucleus send non-canonical, widespread projections to motor and non-motor areas (*20*). Our data suggest that iCNNs control the onset of dystonic crisis through their interaction with the CL. To our knowledge, this is the first evidence to support a role for iCNNs in disease, to describe a functional role for their extracerebellar projections outside the inferior olive, and to characterize their interaction with the motor network in disease. This finding of a precise, cell type-specific circuit may hold promise as a therapeutic avenue for intervention in patients with dystonic crisis or other refractory and debilitating forms of dystonia, with our data raising opportunities for further preclinical and clinical investigation. The significance of uncovering the distinct functions of intermingled neurons, like those of the cerebellar nuclei, has been highlighted in other brain areas and diseases. In a mouse model of Parkinson’s disease, Spix et al. 2021 leveraged differences in cell-type specific neurons within the external globus pallidus to develop an optimized protocol for DBS, leading to longer-lasting therapeutic effects when compared to conventional DBS (*22*). Adapting this approach for optimizing DBS directed to iCNNs and its projections in the CL may yield similarly promising therapeutic results for patients with dystonia or other cerebellar-related disorders. Whether iCNNs contribute to other diseases remains an open question, although we anticipate that our results will encourage the further exploration of this possibility and the therapeutic utility of this understudied neuronal population.

We demonstrated that photoinhibition of iCNN projections reduced the amount of time in spontaneous dystonic crisis, and that these effects changed over multiple days, although the mechanistic cause and location(s) of these changes cannot be determined from our present experiments. Previous studies demonstrated that Arch stimulation could be used over multiple days to slow learning, but they were not designed to demonstrate or refute cumulative effects of stimulation (*62*, *63*). However, studies using channelrhodopsin have demonstrated cumulative, multi-day effects of brief daily optogenetic stimulation. Ahmari et al. 2013 found that 5 minutes of ChR2 stimulation each day for 6 days was sufficient to induce persistently increased grooming in a mouse model of OCD. These behavioral changes were accompanied by an increased responsiveness of postsynaptic neurons to light stimulation, indicating that brief daily optogenetic stimulation induced synaptic plasticity (*64*). Brief daily channelrhodopsin stimulation has also been used to cause increasingly severe seizures in a mouse model of epilepsy, which further demonstrates network plasticity can be induced by optogenetic stimulation over days (*65*).

There are multiple, nonexclusive possible locations for plasticity that could underlie the cumulative multi-day effects of Arch stimulation we observed. iCNNs are innervated by excitatory mossy fibers, which transmit sensory information to the cerebellum, and by inhibitory Purkinje cells. Synchronous Purkinje cell activity leads to hyperpolarization, followed by a rebound depolarization and increased firing, in CNNs (*66*). Foundational research on the mechanisms of plasticity in CNNs highlighted the importance of post-inhibitory rebound depolarization in both long-term potentiation and long-term depression of mossy fiber-CNN and Purkinje cell-CNN synapses (*67–70*). Because the offset of Arch stimulation in the superior cerebellar peduncle has the potential to induce rebound depolarization in essentially all iCNNs simultaneously, it could have widespread effects on the strength of both mossy fiber and Purkinje cell synapses throughout the cerebellar nuclei, although the direction of plasticity would likely depend on the activity of each input at the time of Arch offset. Because stronger hyperpolarization leads to stronger rebound depolarization (*71*) and hyperpolarization paired with mossy fiber activity resulted in stronger potentiation than hyperpolarization alone (*70*), potentiated Purkinje cell-CNN and mossy fiber-CNN synapses may have a higher chance of potentiating further at the next offset of Arch stimulation. This could explain how Arch stimulation led to cumulative, multiday effects. However, each of these possibilities would need to be verified experimentally. Future studies could resolve these possibilities and may even identify additional cell types that are key in the expression and/or alleviation of dystonic crisis.

We demonstrated that photoactivating iCNN projections specifically in the CL (in contrast to the widespread iCNN projections in the superior cerebellar peduncle) modestly but significantly induced dystonic crisis. This indicates that the iCNN to CL projections are involved in the development of dystonic crisis, without excluding a role for the other iCNN projections. It is possible that the relatively small increase in time spent in dystonic crisis was due to insufficient illumination, as we used an LED rather than a laser in these experiments. Additionally, the greater dorsal-ventral length of the CL results in less intense illumination for the deepest projections, compared to those in the superior cerebellar peduncle. Finally, it is likely that other targets of the iCNN projections may be involved in the development of dystonic crisis, and that the CL represents only one node in this network.

We also found that DBS of the CL alleviated dystonic crises that were strongly induced by broader photoactivation of iCNN axons in the superior cerebellar peduncle. While this result suggests that both iCNNs and CL neurons interact to control the onset of dystonic crises, it does not reveal the exact nature of how this interaction occurs. Whether this therapeutic effect relies solely on the iCNN-CL projections, if the modulation of additional inputs to the CL by DBS is essential, how iCNN activity modulates CL activity on a moment-to-moment basis during crisis induction, and how CL activity may further influence other connected motor structures beyond the basal ganglia are all open questions. Elucidating these features may pinpoint the complete repertoire of neural substrates in the motor network that collaborate to induce dystonic crises and how they interact to drive the condition. Answering these questions may reveal critical disease-modifying therapeutic strategies.

Similar to the effects of photoinhibition of iCNN projections, we demonstrated that DBS of the CL reduced the amount of time in induced dystonic crisis, and that these effects changed over multiple days. These improvements could be supported by changes at multiple locations, as DBS can exert several effects simultaneously. These include inducing synaptic vesicle release, activation of efferent axons, and antidromic activation of afferent axons, which can lead to orthodromic action potentials in collaterals of stimulated afferents (*72*). As we found that activating iCNNs induced dystonic crises and inhibiting iCNNs reduced them, the reduction in dystonic crises we observed during CL DBS is consistent with an increase in CL activity (efferent activation). Other groups have shown both immediate (*73*, *74*) and multi-day (*74*, *75*) behavioral effects of electrical stimulation of the intralaminar thalamus (which includes the CL) in rats, and these behavioral changes were accompanied by downstream cellular and molecular effects. Tsai et al. 2000 found that one 30-minute session of intralaminar stimulation normalized the ability of rats injected with β-amyloid to learn the Morris Water Maze over the course of 5 days, and reversed β-amyloid-induced dendritic spine loss and low expression of PSD95 in the cortex and hippocampus (*75*). Shirvalkar et al. 2006 found that 30 minutes of central thalamus stimulation during novel object recognition improved performance, that performance increased for 3 days with repeated stimulation, and that stimulation increased hippocampal and cortical expression of immediate early genes *c-fos* and *zif268* (*74*). These studies suggest that DBS in the CL may lead to molecular and structural changes that support plasticity in the cortex, which could explain the increasing improvement we observed over days of DBS. Evidence of network-wide DBS-induced plasticity has also been identified in dystonic hamsters, where 11 days of DBS in the entopeduncular nucleus (rodent homologue of the globus pallidus internus) normalized firing rates and spike amplitudes in cerebellar neurons and altered the firing properties of thalamic and motor cortical neurons (*76*, *77*). These studies indicate that DBS can induce plasticity that affects neurons throughout the motor network, both in structures monosynaptically connected to the DBS target, as well as those more distantly connected.

Alternatively, the therapeutic effect of DBS in the CL in our mice may be mediated by antidromic activation of iCNN axons. Antidromic activation of hyperdirect pathway axons by subthalamic DBS is thought to be important in disrupting pathological cortical oscillations in Parkinson’s disease (*78*). We have previously identified pathological firing patterns in CNNs in our *Ptf1a^Cre^;Vglut2^fx/fx^*mice, and optogenetically replicating these patterns in behaviorally normal mice induces dystonic phenotypes (*30*, *79*). CL DBS may drive antidromic action potentials in iCNN axons (potentially including collaterals) to create an “information lesion” that masks pathological activity propagating to downstream targets (*80*). However, whether and the extent to which these changes occur in our mice and their relevance to the motor improvements we observed remains to be tested. It would also be interesting to test whether the same therapeutic mechanisms underlying DBS-CL may be shared by photoinhibition of the iCNNs as observed in our study. For example, several studies have shown that in some DBS targets, DBS functions by silencing the target neuronal populations and inducing plasticity (*81*), which may be functionally similar to optogenetic silencing. Such studies may further clarify the therapeutic potential of the iCNNs and their downstream targets, such as the CL.

One type of plasticity that is particularly relevant to DBS is habituation (sometimes known as tolerance), where the effects of continuous stimulation diminish over time and an increase in stimulation intensity is required to maintain stimulation efficacy (*82*, *83*). In our previous study, the same intensity of cerebellar DBS consistently reduced dystonic symptoms over a period of 5 days (*30*). In the present study, we found little evidence of habituation to photostimulation, photoinhibition, or DBS over the course of 3 days.

It is important to note that although we observed cumulative, multi-day effects of photoinhibtion and DBS, we did not test whether these changes would be maintained in the absence of continuing stimulation. In patients with dystonia treated with DBS, improvement often develops slowly over weeks to months, but symptoms return rapidly (within hours to days) if stimulation is turned off (*42*, *84*, *85*). Whether our mice would demonstrate a similar regression in symptoms remains to be systematically tested. Our goal here was to determine whether iCNNs were mediators of dystonic crisis and whether they could be therapeutically targeted.

For decades in clinical research and practice, dystonia severity has been classified using standardized but qualitative scales such as the Burke-Fahn-Marsden Dystonia Rating Scale (*86*), the Global Dystonia Rating Scale (*87*), or the Toronto Western Spasmodic Torticollis Rating Scale (*88*), among others (*89–91*). Similarly, pre-clincial research using rodent models has relied on qualitative scales to quantify dystonia (*19*, *92*). Automated quantification of dystonia in humans is an area of active research (*93–96*), although not yet in clinical use. Likewise, automated assessment of dystonia using kinematic tracking in rodents has been used in some studies (*97*, *98*), but this requires tail suspension, which simultaneously induces dystonic behavior and ensures that all points are visible through the duration of the video. Because we wanted to identify neural populations that could induce dystonic crisis in the absence of acute stress, video analysis of dystonia during tail suspension was not compatible with our study. Automated analysis of dystonia may increase the precision of future studies, but rating scales are as yet the standard in pre-clinical and clinical research. In addition to analyzing behavioral symptoms of dystonic crisis, future studies in mice could incorporate measurements of muscle health and breathing pattern, as these are sometimes compromised in dystonic crises in human patients.

Dystonia produced by our genetic manipulation presents with a range of severity from mouse to mouse, with differences in phenotype extending to the time of day examined, which is similar to the incomplete penetrance and variable presentation of dystonia-linked genes observed in humans (*35*, *36*). For example, even family members with the same *DYT1* mutation may present significantly different dystonia symptoms and severities (*36*). Future studies probing the cause of symptom variability in our mice could further clarify circuits and cell types that contribute to dystonic crisis. In our experiments to determine whether optogenetic inhibition of the iCNN reduces the time spent in dystonic crisis, we intentionally selected animals that exhibited severe dystonic crises more frequently. This selection was intended to avoid a potential floor effect (e.g., where mice are so rarely in dystonic crisis that we cannot detect a decrease in the frequency of dystonic crisis). We did not analyze the spontaneous (without photostimulation) severity of mice in our DBS experiments, but our results show that photostimulation of the superior cerebellar peduncle caused these mice to spend >90% of the time in dystonic crisis, similar to what we observed in our original experiments (**Figure 2F**). Although our current experiments cannot address why baseline severity differs between mice or how mice with different baseline severities responded differently to interventions, they were designed to test the hypothesis that iCNNs are key mediators of dystonic crisis. Importantly, the variability we observe before iCNN stimulation essentially disappears during stimulation (all mice experience near-constant dystonic crisis during stimulation), highlighting the effectiveness of iCNN stimulation to induce dystonic crisis, regardless of baseline symptom severity. Future studies determining whether animals respond differently to optogenetic manipulation and DBS based on baseline symptom severity could further clarify the role of iCNNs and the CL in dystonia outside of dystonic crisis.

Our study utilizes a unique intersectional genetic approach leveraging the developmental, *Ptf1a*-dependent expression of cre, which allowed access to iCNNs in all cerebellar nuclei. This may explain, at least in part, why our anatomical results may appear inconsistent with a recent study. Unlike our study, Judd et al. 2021 did not find that iCNNs project to the CL, which may be due to their viral injection strategy of cre-dependent fluorescent proteins in Vgat-cre mice (*20*). Specifically, the potential iCNNs that could be labeled with this technique were restricted primarily to the injected region (the anterior interposed cerebellar nucleus, a portion of one of three cerebellar nuclei) and dependent on the variability of targeting and viral expression, which may have not included those iCNNs that do project to the CL. In fact, we found CL-projecting iCNNs in all three cerebellar nuclei (**Supplemental Figure 5**).

Our approach of leveraging the *Ptf1a*-specific expression of opsins in iCNNs and the anatomical restriction of their projections in the superior cerebellar peduncle provided an opportunity to selectively modulate their activity while avoiding unwanted photomodulation of nearby regions. While the robust ChR2-YFP expression in the targeted region of the superior cerebellar peduncle validated our approach, we cannot exclude the possibility that other observable nearby Ptf1a-expressing axons in the Kolliker-Fuse nucleus (KF), supratrigeminal nucleus (Su5), and motor trigeminal nucleus (Mo5) may have contributed to the optogenetically induced behaviors we observed (**Supplemental Figure 2A’**). Similarly, *Ptf1a*-expressing projections from Purkinje cells in the parabrachial nucleus, which predominantly target areas around the superior cerebellar peduncle around 5.6 mm posterior to bregma, may have been photomodulated from our approach (*99*). Nonetheless, this is unlikely as the light spread from our setup was calculated to not sufficiently illuminate these regions for photomodulation (see **Methods – optical fiber implantation and stimulation**). Additional evidence that the effects we observed are mediated by stimulating iCNNs rather than other Ptf1a-expressing neurons can be found by examining the connectivity of other potentially-stimulated axons (in the KF, Su5, Mo5 or parabrachial nucleus) and the CL. We demonstrated that optogenetic stimulation of the CL in *Ptf1a^Cre^;Vglut2^fx/fx^;ROSA^lsl-ChR2^*mice moderately but significantly increased the time in dystonic crisis. Additionally, DBS of the CL in these mice significantly reduced the time in dystonic crisis induced by optogenetic stimulation of the superior cerebellar peduncle. However, iCNNs are the only Ptf1a-expressing neurons in or near the superior cerebellar peduncle that have been shown to project to the CL. Although the parabrachial nucleus projects to the thalamic interlaminar nuclei (of which the CL is one), Krout and Loewy 2000 did not find evidence that the parabrachial nucleus projects to the CL (*100*). Further, no studies have shown an anatomical or functional connection between the Su5 or Mo5 and the CL. If photostimulation of the superior cerebellar peduncle was driving dystonic crisis through off-target activation of the KF, Su5, or Mo5, it is difficult to conceive why photostimulation of the CL also drove dystonic crisis, and why DBS of the CL is so effective in reducing dystonic crisis. Moreover, these regions are known to be involved in breathing, the sensorimotor control of the trigeminal nerve, and place preference, which would be unlikely drivers of dystonic crisis. Together, these findings suggest that it is unlikely that the behaviors we reported result from stimulation of Ptf1a-expressing neurons other than iCNNs.

Other brain areas expressing *Ptf1a* were unlikely to be modulated by our manipulation. For example, the Allen Brain Atlas and Human Protein Atlas both show low to no expression of *Ptf1a* mRNA in adults, which cannot be distinguished from unavoidable background staining due to the consistent low amplitude of the signal throughout the brain, as would be expected from noise. It should also be noted that even if the measured expression is true signal rather than background, the expression of *Ptf1a* mRNA may not necessarily correlate with Ptf1a protein expression. Our previous work evaluated the expression of *Ptf1a^Cre^*-driven reporter expression across brain areas (*30*). As already described, *Ptf1a* is developmentally expressed in inhibitory cells of the cerebellum as well as excitatory cells of the inferior olive. It is also expressed in horizontal and amacrine cells of the retina (most of which are GABAergic or glycinergic (*101*, *102*); GABAergic and glycinergic cells of the nucleus of the solitary tract, spinal trigeminal nucleus, principal trigeminal nucleus, and cochlear nucleus; and GABAergic and glycinergic cells of the spinal cord (*103*). As these cells are inhibitory, the removal of *Vglut2* should have little to no direct effect on their function. Additionally, their distance from our optogenetic fibers makes it highly unlikely that they were affected by our photostimulation. Outside of the inferior olive, the only excitatory cells which express *Ptf1a* during development are found in the hypothalamus (*34*). Although these cells may be affected by our genetic manipulation, given the homeostatic role of the hypothalamus, it is unlikely that these cells directly contribute to the specific dystonic phenotypes we observe in our mice. Additional evidence of the cerebellar origin and basis of dystonia in our mice is found in the effectiveness of cerebellar DBS in reducing dystonia (*30*) and our ability to induce dystonic movements in healthy mice by optogenetically replicating the abnormal firing patterns we identified in CNNs in *Ptf1a^Cre^;Vglut2^fx/fx^* mice (*79*).

Although our experiments support a role for the iCNN pathway in controlling the onset of dystonic crisis, it is highly likely that excitatory cerebellar nuclei neurons also contribute to the condition as supported by studies into forms of paroxysmal dystonia. Wu et al. 2025 showed that selective photoactivation of excitatory cerebellar nuclei neurons induced dystonia-like movements in healthy mice, and that selective ablation of these neurons alleviated attacks in mouse models of paroxysmal dystonia (*19*). Other studies similarly supported a contribution for excitatory cerebellar nuclei neurons in paroxysmal dystonia though their optogenetic methods technically did not exclude iCNNs from being manipulated in their studies. While iCNNs project outside the cerebellum and to the cerebellar cortex, they are also known to collateralize widely and form local synaptic connections with excitatory cerebellar nuclei neurons (*17*, *20*, *56*); thus, inhibitory and excitatory cerebellar nuclei neuron activity are tightly linked. Furthermore, Judd et al. 2021 showed that the excitatory and inhibitory cerebellar nuclei share many inputs and extracerebellar outputs (*20*), suggesting that they work synergistically. One possibility is that the excitatory neurons play a major role in the severity of baseline dystonia whereas the inhibitory neurons set the threshold for the dystonic crisis. While we cannot exclude the possibility that our optogenetic manipulation of iCNN activity through ChR2 could have caused backpropagation and influenced excitatory cerebellar nuclei neurons, it still does not discount our finding that direct manipulation of iCNN activity ultimately contributed to the phenotypes we observed in this study, regardless of what other structures may have been involved or required. Future studies are needed to investigate how the excitatory and inhibitory cerebellar nuclei neurons interact to control dystonic crisis and other forms of dystonia, which may reveal population-specific strategies for designing therapeutic interventions to address the broader symptomology that collectively impact patients with dystonia.

## Materials and Methods

### Clinical methods

We performed a retrospective chart review of patients admitted to Texas Children’s Hospital for the primary diagnosis of status dystonicus (also termed dystonic crisis) using (ICD9/10 codes) from 2015-2020 under approved IRB (H-48706). Patients were excluded if subsequent clinical diagnostics excluded status dystonicus in favor of an alternative diagnosis (e.g. status epilepticus). Demographics, medical history, and clinical notes were reviewed for all patients and information included etiology, laboratory studies, imaging, treatments, dosing, response, length of stay, and recurrence of symptoms. Efficacy of treatment was a qualitative measure derived from discussion with patient families and providers as documented within clinical notes. Examples of treatment response measures included de-escalation of treatment after initiation or titration of the medication, improvement in pain scores, improvement in overall tone, and reduction of serum CK. Standardized dystonia rating scales are typically not done routinely in the acute inpatient setting, which prevents their use in our cohort and is a noted limitation of using a retrospective chart review.

### Animals

All experiments were performed following the relevant ethical regulations for animal testing. All mice were housed in an AALAS-accredited facility on a 14h/10h light/dark cycle. Mice of both sexes were studied. Husbandry, housing, euthanasia, and experimental guidelines were reviewed and approved by the Institutional Animal Care and Use Committee (IACUC) of Baylor College of Medicine (protocol number: AN-5996).

Dr Chris Wright (Vanderbilt University School of Medicine) kindly provided the *Ptf1a^Cre^* mice(*30*, *33*). We purchased the *Vglut2^floxed^* (*Vglut2^fx^*, #012898) mice (*104*) from The Jackson Laboratory (Bar Harbor, ME, USA) that contained *LoxP* sites surrounding exon 2 of the gene encoding vesicular glutamate transporter 2 (VGLUT2). These two mouse lines were bred out as described previously to generate dystonic *Ptf1a^Cre^;Vglut2^fx/fx^*mice (*30*). To develop dystonic transgenic mice with restricted Sun1 expression, we crossed these *Ptf1a^Cre^;Vglut2^fx/fx^*mice to heterozygous or homozygous *ROSA^lox-stop-lox-Sun1^*λ lines (*B6;129-Gt(ROSA)26Sor^tm5(CAG-Sun1/sfGFP)Nat^/J*, Strain #:021039, Jackson Laboratory, Bar Harbor, ME, USA). To gain light-sensitive control over inhibitory cerebellar nuclei neurons with channelrhodopsin (ChR2) or archaerhodopsin (Arch), we crossed *Ptf1a^Cre^;Vglut2^fx/fx^* or *Ptf1a^Cre^* males to either *Ai32 ROSA^lox-stop-lox-ChR2-EYFP^* (*B6;129-Gt(ROSA)26Sor^tm32(CAG-COP4*H134R/EYFP)Hze^/J*, Strain #:024109, Jackson Laboratory, Bar Harbor, ME) or *Ai40D* (*Gt(ROSA)26Sor* ^tm40.1(CAG-aop3/GFP)Hze/J^, Strain #:021188, Jackson Laboratory, Bar Harbor, ME) females(*105*). These genotypes of all offspring were confirmed by PCR of digested tissue (tail snips or ear punches) and confirmed by presence of Sun1-GFP, ChR2-EYFP, or Arch-EYFP signal in select neurons (it is important to ensure that cre-dependent expression is not present in all neurons, which would indicate germline transmission). For experimental studies, the day of birth was considered as postnatal day 0 (P0).

### Surgery

Surgery for awake optogenetics was performed as described previously (*106*). In brief, mice were anesthetized with 4% isoflurane in an induction chamber until unresponsive to the toe-pinch reflex. After transferring to a stereotaxic platform (David Kopf Instruments, Tujenga, CA, USA), isoflurane was lowered and maintained at 2% for the duration of surgery. Using aseptic technique and a sterile work field, the mouse was cleaned and prepped and the skin above the skull opened, and a small craniotomy of approximately 1 mm in diameter made above the target region(s). Following the implantation of optical fibers or deep brain stimulation electrodes as described below, all animals were allowed to recover for at least three days to a maximum of one week after surgery. The following surgical techniques were either employed as individual experiments or combined depending on the requirements of the experiment.

### Optical fiber implantation and stimulation

Optical fiber implant surgeries began as described above. One (for unilateral) or two (for bilateral) small craniotomies of about 1mm were performed dorsal to the target region through which optical fibers (Thorlabs, Newton, NJ, USA; #FT200UMT) were lowered into the target region: the superior cerebellar peduncle (5.02 mm posterior and ±1.25 mm lateral to Bregma, 2.5 mm ventral from the brain surface). Optical fibers had been previously glued into ceramic ferrules (Thorlabs, Newton, NJ, USA; #CFLC230-10), polished (Thorlabs, Newton, NJ, USA; #LF5P, #LF3P, #LF1P, #LF03P, #LFCF), epoxied to each other at a set distance using Bondic, (a UV light-activated bonding agent; Bondic, Niagara Falls, NY, USA), and placed inside ceramic mating sleeves (Thorlabs, Newton, NJ, USA; #ADAL 1–5) prior to implantation. The optical fibers and 1-2 metal rods, used for holding the mouse’s head stable while connecting the optical fiber patch cables, were affixed to the skull with C&B Metabond (Parkell, Inc., Edgewood, NY, USA, SKU: S380) and Teets ‘Cold Cure’ Dental Cement (A-M Systems, LLC, Carlsborg, WA, USA, Catalog #525000 and #526000).

Stimulation patterns were programmed and recorded using Spike2 software and delivered using a CED Power1401 data acquisition interface (CED, Cambridge, UK) to control a 470 nm LED (Thorlabs, Newton, NJ, USA; #M470F4) or a 561 nm laser (CrystaLaser, Reno, NV, USA; # CL561-050). Maximum LED power at the end of the implanted fiber was measured to be ∼3.0 mW and stimulation consisted of 50 Hz square pulses of blue 470 nm light. Maximum laser power at the end of the implanted fiber was measured to be ∼10 mW and stimulation consisted of a continuous pulse of green 561 nm light. The light powers were calculated to be capable of driving neurons at a distance of about 0.2 mm away or 0.4 mm away from the fiber tip for 470 nm LED light and 561 nm laser light, respectively, using the threshold of 1 mW/mm^2^ (*107*).

### Tracer injections

This procedure for tracer injections has been described in detail previously (*108*). Briefly, prior to surgery, the glass micropipettes (Borosilicate Standard Wall without Filament glass capillaries, 1.5 mm o.d. × 0.86 mm i.d. × 75 mm length) were prepared using a micropipette puller with the heat level set at 60 on the Step 1 setting of the Narishige PC-10 Puller and with two heavy weights and one light weight attached. Prior to use, the micropipette tips were clipped using forceps to ensure dispensability of neural tracer, the micropipette was attached to the Nanoject II, and ∼1.0 microliters of BDA 3k were loaded into the micropipette. Then, the surgery proceeded as described above for the optical fiber implantation. Except, instead of optical fibers, the filled micropipette was lowered slowly into the stereotaxic coordinates for the centrolateral nucleus of the thalamus (1.50 mm posterior and ±0.80 mm lateral to Bregma, 3.25 mm ventral from the brain surface). The Nanoject II was set to inject the smallest possible amount at the slowest available speed, and injections were given at 10 second intervals until ∼75 nL of BDA 3K were injected. After waiting five minutes, the micropipette was slowly taken out of the brain. The procedure was repeated for the contralateral site. After injections, the craniotomies were covered with antibiotic and the skull and skin were sealed with C&B Metabond (Parkell, Inc., Edgewood, NY, USA, SKU: S380) and Teets ‘Cold Cure’ Dental Cement (A-M Systems, LLC, Carlsborg, WA, USA, Catalog #525000 and #526000). Mice were transcardially perfused 9-12 days after the tracer injection procedure to allow the BDA 3K to penetrate neurons and travel retrogradely into the soma of input neurons (*52*, *53*).

### DBS electrode implantation and stimulation

DBS electrode implant surgeries were performed exactly as the optical fiber implantation surgeries. Except, instead of optical fibers, custom 50 mm twisted bipolar Tungsten DBS electrodes were used (PlasticsOne, Roanoke, VA, USA; #8IMS303T3B01) and targeted to the centrolateral nucleus of the thalamus (1.50 mm posterior and ±0.78 mm lateral to Bregma, 2.55 mm ventral from the brain surface). For bilateral implantation, the two electrodes were fixed together using Bondic, a UV light-activated bonding agent (Bondic, Niagara Falls, NY, USA). The electrodes and 1-2 metal rods, used for holding the mouse’s head stable while connecting the DBS cables, were secured to the head with C&B Metabond (Parkell, Inc., Edgewood, NY, USA, SKU: S380) and Teets ‘Cold Cure’ Dental Cement (A-M Systems, LLC, Carlsborg, WA, USA, Catalog #525000 and #526000).

For DBS of the centrolateral nucleus of the thalamus, we used a Multichannel Systems Pulse generator (AMPI, Jerusalem, Israel) programmed to deliver 30 μA, 60 μs biphasic square pulses at 130 Hz.

### Behavioral analyses

The *Ptf1a^Cre^;Vglut2^fx/fx^* mutant mice exhibit dystonic symptoms of mild-to-moderate twisting postures in the limbs, trunk, head, and/or tail interrupted by random and intermittent attacks of generalized dystonia in several body parts reminiscent of dystonic crises (*30*). Similar to human patients in which the same underlying genetic mutation may result in a spectrum of dystonic symptoms across patients (*35*, *36*), the severity of the underlying dystonic symptoms in the *Ptf1a^Cre^;Vglut2^fx/fx^* mouse can vary between animals. Spontaneous and optogenetically-induced dystonic crises were marked by prolonged and severe dystonic posturing including extended limbs, rigidity throughout the body, stiff tail, splayed digits, toppling over, and severely impaired ambulation for at least three seconds. Optogenetically-induced dystonic crises persisted throughout the length of the optogenetic stimulation, except during or following DBS of the centrolateral nucleus of the thalamus.

The behavior of experimental mice in an open field was video recorded and video data from before and during stimulation were analyzed. For experiments involving photoinhibition (**Figure 3B**) and DBS of the CL (**Figure 4G**), three trials were given each day for four consecutive days. To detect potential changes in the incidence of dystonic crises, we measured the percentage of time that a mouse spent in a dystonic crisis during each period (before vs during stimulation), calculated the average for each period on a given day if applicable, and made statistical comparisons using the Wilcoxon matched-pairs signed rank test. Throughout this study, we define a dystonic crisis as observable dystonic muscle contractions or postures in at least two body parts in addition to severely limited ambulation for at least 3 seconds, a sufficient time to confidently determine that ambulation was limited (rather than due to resting) and a result of abnormal muscle activity.

For bilateral photoactivation of the superior cerebellar peduncle in *Ptf1a^Cre^;Vglut2^fx/fx^;ROSA^lsl-Chr2/EYFP^* mice, two-minute periods were video recorded before and during stimulation. For bilateral photoinhibition of the superior cerebellar peduncle in *Ptf1a^Cre^;Vglut2^fx/fx^;ROSA^lsl-Arch/EYFP^*mice, behavioral video data was collected over four consecutive days. To establish a baseline for how much time the mice exhibit dystonic crises over a 35-minute session without optogenetic manipulation, we included a sham-stimulation day on Day 0, in which the mice were connected to the optic fibers throughout the 35-minute session as on subsequent days, but no light stimulation was administered (**Figure 3B**). To test whether photostimulation influenced the percentage of time spent in dystonic crisis, video was recorded for five minutes before and five minutes during stimulation. To increase confidence that any behavioral change was a result of optogenetic manipulation, we repeated the stimulation paradigm three times on each day, resulting in a 35-minute session for each day (**Figure 3B**).

For unilateral photoactivation of the superior cerebellar peduncle and bilateral DBS of the CL in *Ptf1a^Cre^;Vglut2^fx/fx^;ROSA^lsl-Chr2/EYFP^*mice, we collected behavioral video data over four consecutive days. Each day consisted of an 86-minute session divided into five distinct periods: photobaseline, DBS period, immediately post-DBS, 30 minutes post-DBS, and 1 hour post-DBS (**Figure 4G**). During each distinct period, a 2-minute stimulation of blue light (470 nm light; 50 Hz square pulses; ≥ 1mW at fiber tip) was administered followed by 2 minutes of no light stimulation. This paradigm was repeated three times for each distinct time period. The percentage of time spent in dystonic crisis was quantified for each of the three light pulses, and the average of these values was analyzed. To establish a baseline for how much time the mice spend in a dystonic crisis induced by iCNN photoactivation, we included a sham-DBS day on Day 0, in which the mice were connected to the stimulus generator throughout the 86-minute session as on subsequent days, but no DBS was administered (**Figure 4G**). DBS was administered on Days 1-3. DBS was only performed in the DBS period for 10 minutes (130 Hz, 60µs, ±30µA), starting with the onset of the first light pulse and ending with the offset of the third light pulse.

### Tissue processing and immunohistochemistry

Perfusion and tissue fixation were performed as previously described (*109*). In brief, mice were anesthetized with Avertin (2, 2, 2-Tribromoethanol, Sigma-Aldrich, St. Louis, MO, USA; #T48402) via intraperitoneal injection. Once mice were deeply anesthetized, a whole-body perfusion was performed first with ice-chilled 0.1M phosphate-buffered saline (PBS; pH 7.4), then with ice-chilled 4% paraformaldehyde (4% PFA) diluted in PBS. The brain was then dissected out and placed in 4% PFA for 24 to 48 hr for post-fixation at 4°C. Cryoprotection was then performed by placing the tissue in stepwise sucrose dilutions, first in 15% sucrose in PBS followed by 30% sucrose in PBS. Brains were stored at 4°C during stepwise sucrose incubation steps. After cryoprotection, the tissue was embedded in Tissue-Tek O.C.T. Compound (Sakura, Torrance, CA, USA) and frozen at −80°C. For immunohistochemistry, free-floating sections cut at 40um on the cryostat were incubated in antibodies in a solution of 10% normal goat or donkey serum and 0.01% Tween-20. For all stains, sections were blocked for 2 h with the blocking solution at room temperature and then incubated in primary antibodies overnight at room temperature. Sections were then washed with PBS 3 times for 5 min each and then incubated in secondary antibodies for 2 h. Sections were washed again 3 times in PBS for 5 min each and mounted onto slides. Slides with fluorescent signal were immediately cover-slipped using FluoroGel with Dapi as a medium. Antibodies used for immunohistochemistry (primary antibodies) were: GFP to visualize ChR2 and Sun1-GFP (Ch, Abcam, 1:1000), NeuroTrace fluorescent Nissl 530/615 to visualize neurons (Molecular Probes Inc, Eugene, OR, USA, #N21482; 1:1500), and Streptavidin Alexa Fluor 647 to visualize neurons filled with BDA 3K (Thermo Fisher Scientific, Eugene, OR, USA, #S21374); Secondary antibodies were: donkey anti-chicken secondary antibodies conjugated to 488 fluorophores (Invitrogen) diluted to 1:1500.

### *In situ* hybridization (ISH)

ISH was performed by the RNA ISH Core at Baylor College of Medicine using an automated robotic platform as described previously (*110*). ISH was used to visualize *Vglut2*, *Vgat,* and *YFP* expression in unfixed, fresh frozen tissue cut in 25 μm-thick coronal brain sections. Digoxigenin (DIG)-labeled mRNA antisense probes against *Vglut2*, *Vgat,* and *YFP* were generated using reverse-transcribed mouse cDNA as a template and an RNA DIG-labeling kit from Roche. Primer and probe sequences for the *Vglut2, Vgat*, and *YFP* probes are available on the Allen Brain Atlas website (http://www.brain-map.org).

### Imaging of immunostained tissue sections

Photomicrographs of stained tissue sections were captured using either a Zeiss Axio Imager.M2 microscope equipped with Zeiss AxioCam MRm and MRc5 cameras (Zeiss, Oberkochen, Germany) or a Leica DM4000 B LED microscope equipped with Leica DFC365 FX and Leica DMC 2900 cameras (Leica Microsystems Inc, Wetzlar, Germany). Zeiss Zen software was used for image acquisition from the Zeiss microscope. Leica Application Suite X (LAS X) software was used for image acquisition from the Leica microscope. Images were corrected for brightness and contrast using Adobe Photoshop CS5 (Adobe Systems, San Jose, CA, USA) for figure preparation. Schematics were made in Adobe Illustrator CC.

### Data analysis and statistics

Data are presented as mean±s.e.m. and were analyzed with a Wilcoxon matched-pairs signed rank test (non-normally distributed data with unequal variance). For all statistical tests, *P*<0.05 was considered as statistically significant. All statistical analyses were performed using GraphPad Prism (GraphPad Software, La Jolla, CA, USA) software.

### Data availability

The data that support the findings of this study are available from the corresponding author upon request.

## Funding

Baylor College of Medicine

Texas Children’s Hospital

The Jan and Dan Duncan Neurological Research Institute (Texas Children’s Hospital Duncan NRI)

The National Institute of Neurological Disorders and Stroke grants R01NS119301 and R01NS127435 (RVS)

The National Institute of Neurological Disorders and Stroke grant K08NS121600 (JSG)

Eunice Kennedy Shriver National Institute of Child Health and Human Development of the National Institutes of Health under Award Number P50HD103555 for use of the Cell and Tissue Pathogenesis Core and In Situ Hybridization Core (the BCM IDDRC) (RVS)

The Ting Tsung and Wei Fong Chao Foundation (RVS)

The content is solely the responsibility of the authors and does not necessarily represent the official views of the National Center for Research Resources or the National Institutes of Health.

### Contributions

Conceptualization: AGRH, and RVS

Methodology: AGRH, MPD, JSG, MH, and RVS

Software: AGRH, MPD

Validation: AGRH, MPD

Formal analyses: AGRH, MPD

Investigation: AGRH, MPD, JSG, JEA, KAC, MH, and RVS

Data curation: AGRH, MPD, JSG, KAC, MH

Writing—original draft: AGRH, MPD, RVS

Writing—review & editing: AGRH, MPD, JSG, JEA, KAC, MH, and RVS

Visualization: AGRH, MPD, JSG, KAC, MH, and RVS

Supervision: RVS

Project administration: AGRH, MPD, JSG, KAC, MH, and RVS

Funding acquisition: JSG and RVS

## Data and materials availability

All data, code, and materials are available in the main text, supplementary materials, or available upon request to the corresponding author.

---

**Supplemental Figure 1:**
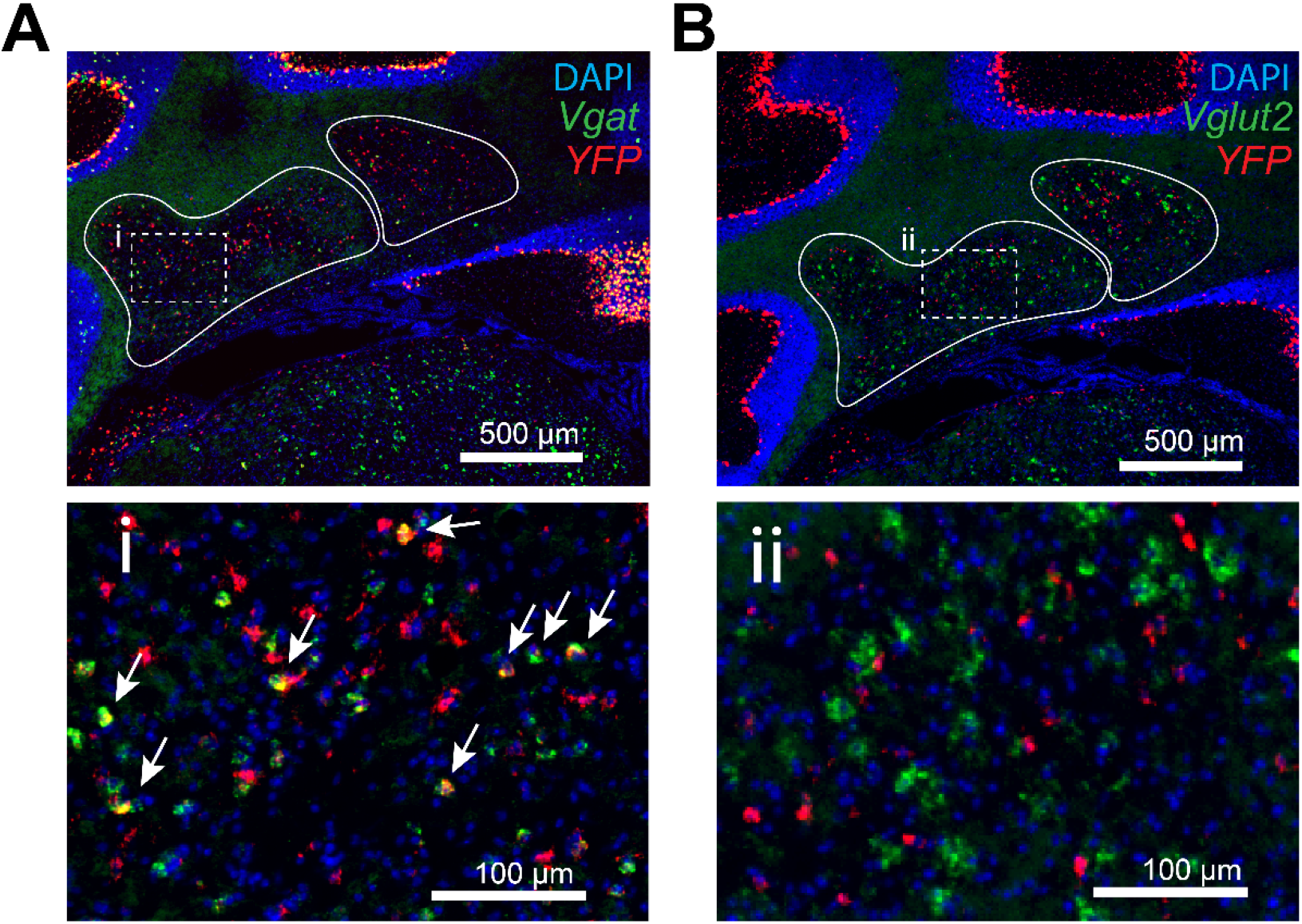
Ptf1a-cre induced expression of cre-dependent ChR2-YFP in inhibitory cerebellar nuclei neurons, but not excitatory cerebellar nuclei neurons, in *Ptf1a^Cre^;Vglut2^fx/fx^;ROSA^lsl-Chr2/EYFP^* mice. (**A**) *In situ* hybridization in a coronal section showing mRNA expression *Vgat* and *YFP* as well as DAPI staining (n=3 animals; 3 sections per animal). The cerebellar nuclei are outlined in white. Inset (i) is a higher magnification image from (**A**) with white arrows marking cerebellar nuclei neurons that were co-labeled with *Vgat* and *YFP* probes. (**B**) *In situ* hybridization in a coronal section showing mRNA expression *Vglut2* and *YFP* as well as DAPI, with the cerebellar nuclei outlined in white (n=3 animals; 3 sections per animal). Inset (ii) is a higher magnification image of (**B**) showing that there was no co-labeling of *Vgut2* and *YFP*.

**Supplemental Figure 2:**
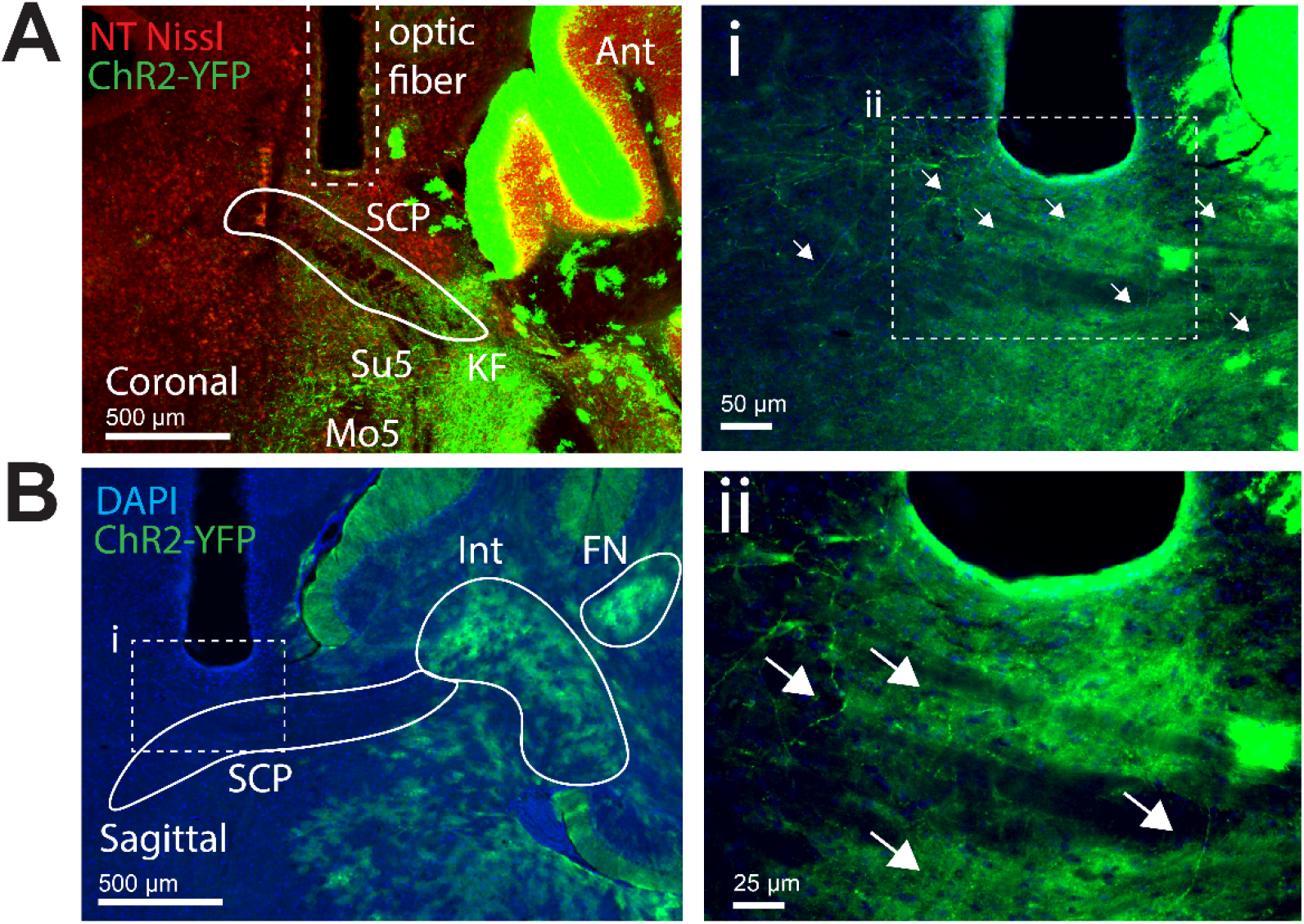
Anatomical verification of optical fibers over the superior cerebellar peduncle (SCP), which houses extracerebellar projections from inhibitory cerebellar nuclei neurons as labeled with ChR2-YFP in *Ptf1a^Cre^;Vglut2^fx/fx^;ROSA^lsl-Chr2/EYFP^*mice. (**A**) Immunohistochemical stain of a coronal section verifying the stereotaxic location of the SCP used for targeting the optical fibers (n=9 animals, 3 sections per animal). Note the punctate labeling of axonal projections expressing ChR2-YFP in the SCP. (**B**) Immunohistochemical stain of a sagittal section verifying the stereotaxic location of the SCP used for targeting the optical fibers (n=3 animals, 3 sections per animal). Insets (i and ii) show higher magnification of the SCP from (**B**), highlighting the expression of ChR2-YFP in axonal projections with white arrows. SCP, superior cerebellar peduncle; Ant, anterior cerebellar lobe; KF, Kolliker-Fuse nucleus; Su5, supratrigeminal nucleus; Mo5, trigeminal motor nucleus; Int, interposed cerebellar nucleus; FN, fastigial cerebellar nucleus.

**Supplemental Figure 3:**
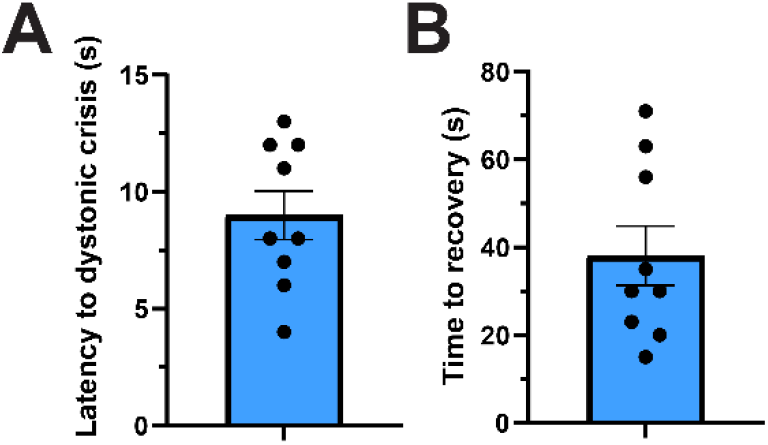
Photoactivation of the inhibitory cerebellar nuclei neuron pathway through the superior cerebellar peduncle induced dystonic crises that occurred within a short latency from light onset and that persisted many seconds after stimulation ends in *Ptf1a^Cre^;Vglut2^fx/fx^;ROSA^lsl-Chr2/EYFP^*mice. (**A**) Dystonic crises were induced at around 10 seconds from light onset. (**B**) After the light stimulation ends, the induced dystonic crisis persisted for around 40 seconds. n=9 animals.

**Supplemental Figure 4:**
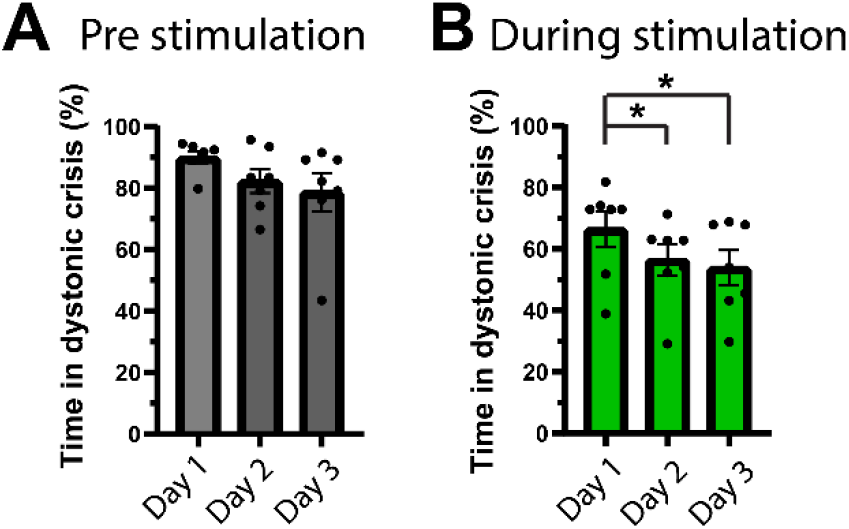
Photostimulation over several days resulted in a reduction in the time spent in dystonic crisis. (**A**) For the pre stimulation periods when comparing Day 1 (the first day of stimulation) against Day 2 (p=0.0625) and Day 3 (p=0.0781), there was a trend toward a reduction in dystonic crisis, but it was not statistically significant. (**B**) For the during stimulation periods, there was a significant reduction in dystonic crises when comparing Day 1 to Day 2 (p=0.0312) and Day 3 (p=0.0156). For all statistics, n=7 animals, Wilcoxon matched-pairs signed rank test, two tailed.

**Supplemental Figure 5:**
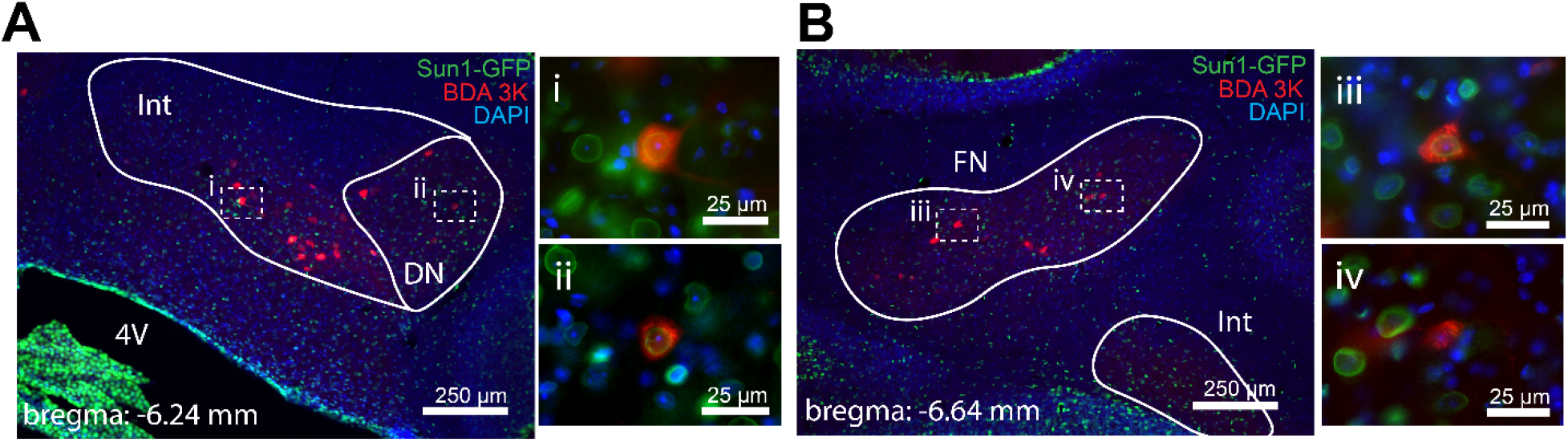
Inhibitory cerebellar nuclei neurons from all three major cerebellar nuclei (fastigial, interposed, dentate) project to the centrolateral nucleus of the thalamus (n=4 animals, 3 sections per animal). (**A**) Coronal section showing that neurons from the interposed and dentate cerebellar nucleus were filled with BDA 3K following injection of BDA 3K, a predominantly retrograde tracer, in the centrolateral nucleus of the thalamus. Insets (i) and (ii) are higher magnification images of (**A**) showing the co-labeling of DAPI, Sun1-GFP, and BDA 3K in the interposed and dentate cerebellar nucleus, respectively. (**B**) Coronal section showing that neurons from the fastigial cerebellar nucleus were filled with BDA 3K following injection of BDA 3K, a predominantly retrograde tracer, in the centrolateral nucleus of the thalamus. Inset (iii) is a higher magnification image of (**B**) showing a co-labeling of DAPI, Sun1-GFP, and BDA 3K. Inset (iv) is a higher magnification image of (**B**) showing co-labeling of DAPI and BDA 3K, but not Sun1-GFP. Int, interposed cerebellar nucleus; DN, dentate cerebellar nucleus; FN, fastigial cerebellar nucleus; 4V, fourth ventricle.

**Supplemental Figure 6:**
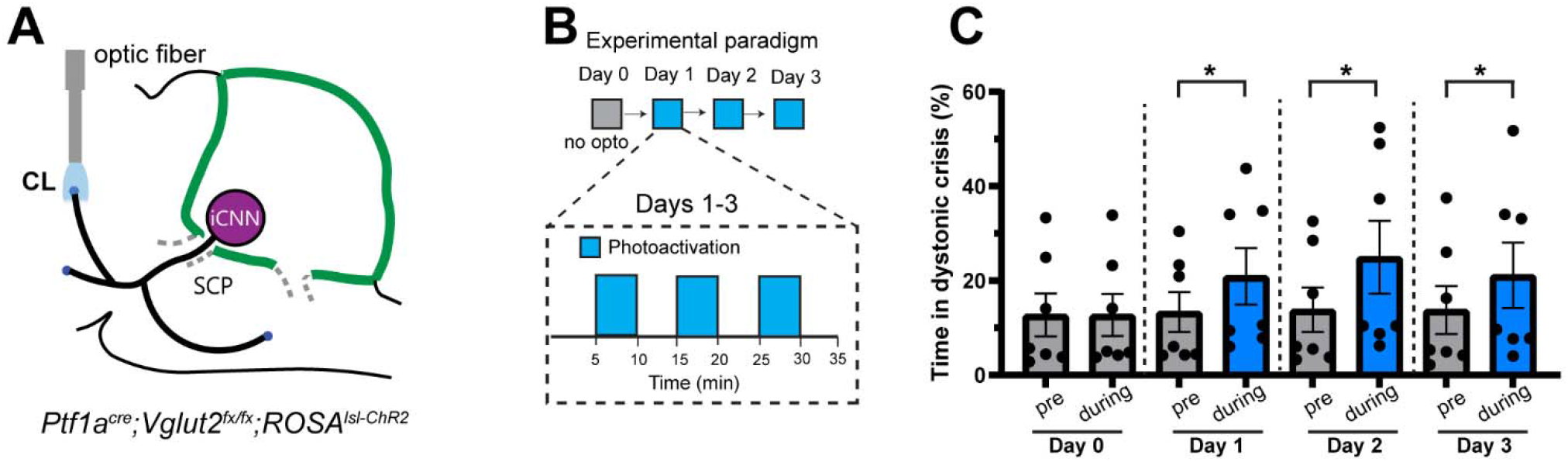
Photostimulation of inhibitory cerebellar nuclei neuron projections in the centrolateral nucleus of the thalamus modestly increases time spent in dystonic crisis in *Ptf1a^Cre^;Vglut2^fx/fx^;ROSA^lsl-ChR2^*mice. (**A**) Schematic showing the selective photoactivation of iCNN terminals in the CL by implanting optical fibers over the CL in *Ptf1a^Cre^;Vglut2^fx/fx^;ROSA^lsl-ChR2^*mice. (**B**) Schematic showing the four-day experimental paradigm. (**C**) Photoactivation of iCNN terminals in the CL modestly but significantly increased the percentage of time spent in crisis when comparing pre versus during photostimulation on Day 1 (p=0.0156), Day 2 (p=0.0156), and Day 3 (p=0.0156), but not on Day 0 (sham stimulation, p=0.8125). For all statistics: n=7 animals, Wilcoxon matched-pairs signed rank test, two-tailed, error bars are defined as standard error of the mean.

**Supplemental Figure 7:**
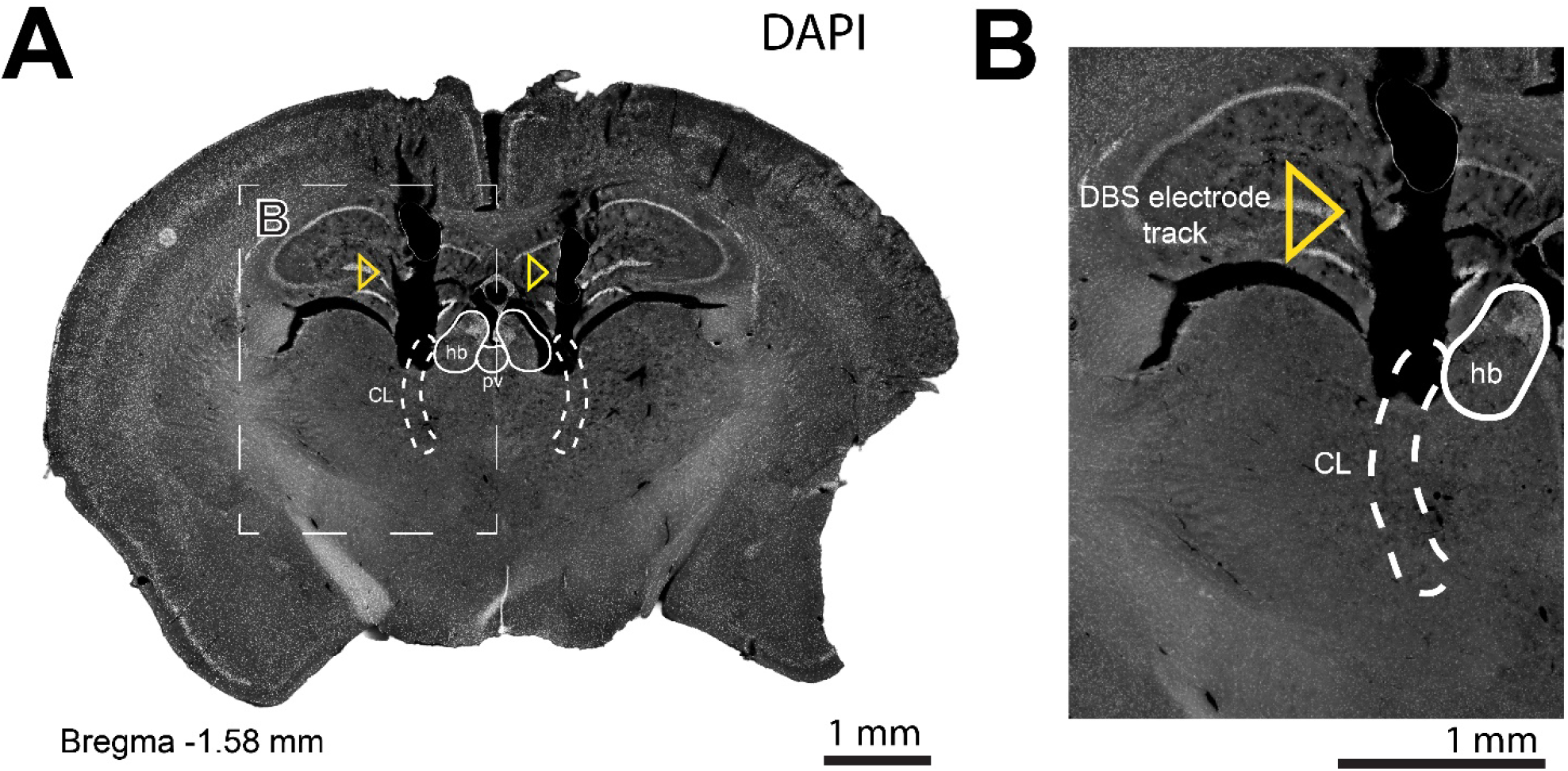
Anatomical verification of the surgical targeting of deep brain stimulation electrodes in the centrolateral nucleus of the thalamus. (**A**) Immunohistochemical staining of DAPI showing the tracks formed by the bilateral implantation of deep brain stimulation electrodes in the centrolateral nucleus of the thalamus (n=6 animals, 3 sections per animal). Yellow triangles mark the tracks formed by the DBS electrodes above and with some penetration into the centrolateral nucleus of the thalamus. (**B**) Inset of A showing a higher magnified image of the DBS electrode track in the centrolateral nucleus of the thalamus. CL, centrolateral nucleus of the thalamus; hb, habenula; pv, paraventricular thalamic nucleus.

**Supplemental Figure 8:**
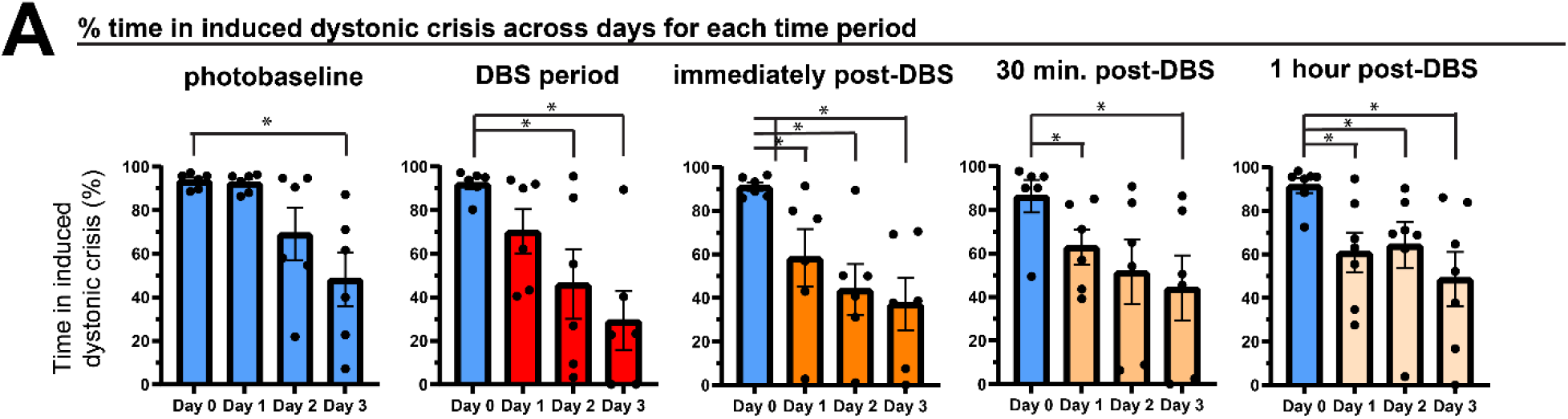
Deep brain stimulation of the centrolateral nucleus of the thalamus over several days reduced the time in dystonic crises induced by photoactivation of the inhibitory cerebellar nuclei neuron pathway. (**A**) Comparing the induced dystonic crises during photobaseline on Day 0 against Day 3 revealed a significant reduction in dystonic crises (p=0.0312). Photoinduced dystonic crises during DBS on Day 0 were reduced on Day 2 (p=0.0312) and Day 3 (p=0.0312). Photoinduced dystonic crises immediately post-DBS on Day 0 were decreased on Day 1 (p=0.0312), Day 2 (p=0.0312), and Day 3 (p=0.0312). For 30 minutes post-DBS, photoinduced crises were reduced on Day 1 (p=0.0312) and Day 3 (p=0.0312). For 1 hour post-DBS, photoinduced crises decreased on Day 1 (p=0.0156), Day 2 (p=0.0156), and Day 3 (p=0.0156). For all statistics: n=6 animals, Wilcoxon matched-pairs signed rank test, two-tailed, error bars are defined as standard error of the mean.

**<u>Supplementary Movie 1</u>**: Movie of a patient in dystonic crisis at the hospital that was included in the study. The patient exhibits the debilitating dystonic postures characteristic of dystonic crisis including rigidity and stiffness in the neck, arms, fingers, and toes.

**<u>Supplementary Movie 2</u>**: Movie of a *Ptf1a^Cre^;Vglut2^fx/fx^;ROSA^lsl-Chr2/EYFP^*mouse before, during, and after photoactivation of the inhibitory cerebellar nuclei neuron pathway via the superior cerebellar peduncle. A dystonic crisis was induced within several seconds of light onset, and the crisis persisted for many seconds after the end of light stimulation.

**<u>Supplementary Movie 3</u>**: Movie of a *Ptf1a^Cre^;ROSA^lsl-Chr2/EYFP^* mouse before, during, and after photoactivation of the inhibitory cerebellar nuclei neuron pathway via the superior cerebellar peduncle. Although no dystonic crisis was induced, the animal moved abnormally with slowed movement, mild dystonia-like muscle tension, crawling, widened hind legs, dragging of the trunk, clunky locomotion, and wide steps.

**<u>Supplementary Movie 4</u>**: Movie of a *Ptf1a^Cre^;Vglut2^fx/fx^;ROSA^lsl-Arch/EYFP^*mouse before and during photoinhibition of the inhibitory cerebellar nuclei neuron pathway via the superior cerebellar peduncle on Day 1 and Day 3 of photostimulation. The first half of the video consisted of footage from Day 1 of stimulation where there was a notable alleviation of the spontaneous dystonic crisis and restoration of mobility. On Day 3 of stimulation as shown in the second half of the video, the baseline time spent in spontaneous dystonic crisis was slightly less severe than Day 1, and photostimulation further improved the mobility of the dystonic mouse to a level that was better than that seen on Day 1 during photostimulation.

<u>Supplementary Movie 5</u>: Movie of a *Ptf1a^Cre^;Vglut2^fx/fx^;ROSA^lsl-Chr2/EYFP^* mouse on Day 1 and Day 3 of deep brain stimulation (DBS) in the centrolateral nucleus of the thalamus (CL) showing the short-term and long-term therapeutic effects of DBS on dystonic crises induced by iCNN photoactivation. On Day 1, the photobaseline shows that photoactivation of the iCNN pathway via the superior cerebellar peduncle reliably induced a dystonic crisis; however, when DBS of the CL was administered concurrently with the light stimulation in the “during DBS” period, a dystonic crisis was induced only briefly and then the mouse quickly recovered its mobility. On Day 3, the photobaseline period shows that iCNN photoactivation still reliably induced dystonic crisis; however, concurrent DBS of the CL in the “during DBS” period prevented the occurrence of a dystonic crisis induced by iCNN photoactivation.

